# Simultaneous smoothing and detection of topological units of genome organization from sparse chromatin contact count matrices with matrix factorization

**DOI:** 10.1101/2020.08.17.254615

**Authors:** Da-Inn Lee, Sushmita Roy

## Abstract

The three-dimensional (3D) organization of the genome plays a critical role in gene regulation for diverse normal and disease processes. High-throughput chromosome conformation capture (3C) assays, such as Hi-C, SPRITE, GAM, and HiChIP, have revealed higher-order organizational units such as topologically associating domains (TADs), which can shape the regulatory landscape governing downstream phenotypes. Analysis of high-throughput 3C data depends on the sequencing depth, which directly affects the resolution and the sparsity of the generated 3D contact count map. Identification of TADs remains a significant challenge due to the sensitivity of existing methods to resolution and sparsity. Here we present GRiNCH, a novel matrix-factorization-based approach for simultaneous TAD discovery and smoothing of contact count matrices from high-throughput 3C data. GRiNCH TADs are enriched in known architectural proteins and chromatin modification signals and are stable to the resolution, and sparsity of the input data. GRiNCH smoothing improves the recovery of structure and significant interactions from low-depth datasets. Furthermore, enrichment analysis of 746 transcription factor motifs in GRiNCH TADs from developmental time-course and cell-line Hi-C datasets predicted transcription factors with potentially novel genome organization roles. GRiNCH is a broadly applicable tool for the analysis of high throughput 3C datasets from a variety of platforms including SPRITE and HiChIP to understand 3D genome organization in diverse biological contexts.

## Introduction

The three-dimensional (3D) organization of the genome has emerged as an important layer of gene regulation in developmental processes, disease progression, and evolution [1–6]. High-throughput chromosome conformation capture (3C) assays such as Hi-C [7, 8], SPRITE [9], and GAM [6] provide a comprehensive view of 3D organization by measuring interactions among chromosomal regions on a genome-wide scale. High-throughput 3C data captured from diverse biological contexts and processes has led to an improved understanding of DNA packaging in the nucleus, and the dynamics of 3D conformation across developmental stages [10], and between normal and disease cellular states [4,11]. Analysis of such datasets has shown that chromosomal regions preferentially interact with one another, giving rise to higher-order structural units such as chromosomal territories, compartments, and topologically associating domains (TADs) which differ in the size of the structural unit and molecular features associated with the constituent regions. Although the relationship between TADs and changes in gene expression is debated [12–14], these units have been shown to be conserved across species [5, 15] and also associated with developmental [16] and disease processes [11, 17–19]. Accurate identification of TADs is an important goal for linking 3D genome organization to cellular function.

Recently a large number of methods to identify TADs have been developed, utilizing different computational frameworks, such as dynamic programming, [20, 21], community and subgraph detection within networks [20, 22], Gaussian mixture modeling [23, 24], and signal processing approaches [25]. However, comparison of TAD-finding methods [26–28] have found large variability in the definition of TADs and high sensitivity to the resolution (size of the genomic region), sequencing depth, and sparsity of the input data. A lack of a clear definition for a TAD leads to difficulty in downstream interpretation of these structures [29]. To address the sparsity of datasets, different smoothing based approaches have been proposed, e.g. mean filter [30] and Gaussian filter [31]; however, it is unclear whether and to what extent TAD identification can benefit from pre-smoothing the matrices.

Here, we present Graph Regularized Non-negative matrix factorization and Clustering for Hi-C (GRiNCH), a novel matrix-factorization-based method for the analysis of high-throughput 3C datasets. GRiNCH is based on non-negative matrix factorization (NMF), a powerful dimensionality reduction method used to recover interpretable low-dimensional structure from high-dimensional datasets [32–34]. However, a standard application of NMF is not sufficient because of the strong distance dependence found in Hi-C data, that is, regions that are close to each other on the linear genome tend to have more interactions. We employ a graph regularized NMF approach, where the graph captures the distance dependence of contact counts such that the learned lower-dimensional representation is smooth over the graph structure [35]. Furthermore, by exploiting NMF’s matrix completion property, which imputes missing entries of a matrix from the product of the low-dimensional factors, GRiNCH can smooth a sparse input matrix.

We perform a comprehensive comparison of GRiNCH and existing TAD-finding methods using a number of metrics: similarity of interaction profiles of regions belonging to the same TAD, stability to different resolutions and depth of input data, and enrichment of architectural proteins and histone modification known to facilitate or correlate with 3D genome organization. Despite the general trend of trade-off in performance among different criteria, e.g., a high performing method based on enrichment of architectural proteins is not as stable to resolution and depth, GRiNCH consistently ranks among the top across different measures. Furthermore, compared to existing smoothing approaches, GRiNCH-based smoothing of downsampled data leads to the recovery of TADs and significant interactions best in agreement with those from the original high-depth dataset. We apply GRiNCH to Hi-C data from two different developmental time courses; we successfully recapitulate previously identified topological changes around key genes, and predict novel boundary factors that could interact with known architectural proteins to form topological domains. Taken together, GRiNCH is a robust and broadly applicable approach to discover structural units and smooth sparse high-throughput 3C datasets from diverse platforms including Hi-C, SPRITE and HiChIP.

## Results

### GRiNCH, a non-negative matrix factorization-based method for analyzing high-throughput chromosome conformation capture datasets

GRiNCH uses graph-regularized Non-negative Matrix Factorization (NMF) to identify topologically associating domains (TADs) from a high-dimensional 3C count matrix (**Figure 1, Methods**). GRiNCH has several properties that make it attractive for analyzing these count matrices: (1) matrix factorization methods including NMF have a “matrix completion” capability, which can be used to smooth noisy, sparse matrices, (2) the low-dimensional factors provide a clustering of the row and column entities that can be used to define chromosomal structural units, (3) the non-negativity constraint of the factors provide a parts-based representation of the data and is well suited for count datasets (such as Hi-C matrices), and (4) GRiNCH can be applied to any symmetric count matrix measuring chromosomal interactions between genomic loci such as Hi-C, [36], SPRITE [9], and HiChIP [37] datasets.

**Figure 1:**
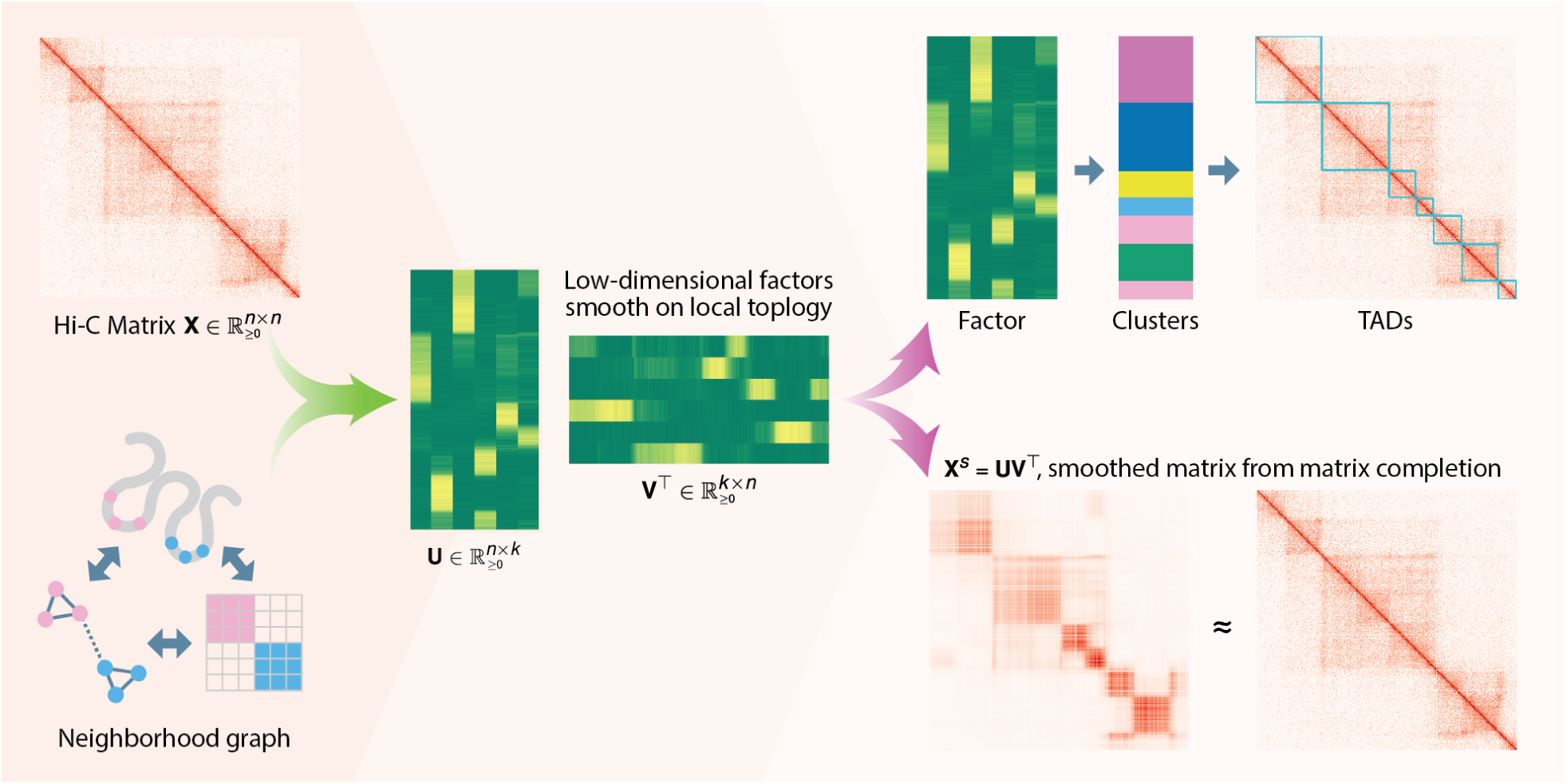
Overview of GRiNCH. GRiNCH applies Non-negative Matrix Factorization (NMF) to a Hi-C or a similar high-throughput 3C matrix to find clusters of densely interacting genomic regions. NMF recovers low-dimensional factors U and V of the input matrix X that can be used to reconstruct the input matrix. As nearby genomic regions tend to interact more with each other, we regularize the factor matrices with a neighborhood graph to encourage neighboring regions to have a similar lower-dimensional representation, and subsequently belong to the same cluster. We cluster the regions by treating one of the factor matrices as a set of latent features and applying *k*-medoids clustering. The clusters represent topological units such as TADs. The factor matrices can be multiplied together to yield a smoothed version of the input matrix which is often sparse and noisy.

For the ease of description, we will consider a Hi-C matrix as the input to GRiNCH. In GRiNCH, the count matrix is approximated by the product of two lower dimensional matrices, U and V, both with dimension *n* × *k*, where *n* is the number of genomic regions in the given chromosome, and *k* is the rank of the lower-dimensional space. Because Hi-C matrices have a strong distance dependence, we use a constrained formulation of NMF, where the columns of the U and V matrices are smooth on a graph of genomic regions (**Figure 1**), such that regions that are connected in the graph have similar sets of values in the lower-dimensional space. The graph in turn captures the distance dependence using a local neighborhood, where two regions *i* and *j* have an edge between them if they are within a particular radius *r* of each other in linear distance along the chromosome. GRiNCH has three parameters, *k*, the rank of the lower dimensional space, *r* to control the size of the neighborhood, and *λ* to control the strength of graph regularization. After factorization, GRiNCH uses chain-constrained *k*-medoids clustering to define clusters of contiguous regions, which we consider as TADs. We probed the impact of the three parameters, *k, r*, and *λ*, on the resulting GRiNCH TADs. We determined that setting *k* to identify TADs of size ≈ 1Mb, with a neighborhood size of *r* = 250kb and a small amount of regularization (*λ* = 1), yields the best results (**Figure S1**). Notably, the regularization yields TADs with higher CTCF enrichment than vanilla matrix factorization without any regularization (i.e. *λ* = 0).

### GRiNCH TADs are high quality and stable to varying resolution and depth of input Hi-C data

To assess the quality of GRiNCH TADs, we considered seven existing TAD identification methods (**Table 1**) and applied them along with GRiNCH to Hi-C data of five different cell lines from Rao et al. [36] for comparison. The quality of a TAD was measured with two internal validation metrics used for cluster evaluation, Davies-Bouldin index (DBI) and Delta Contact Count (DCC), both assessing the similarity of interaction profiles of regions within defined TADs. DBI of a cluster measures how well separated the given cluster is from other clusters; in our case, how distinct each TAD’s interaction count profile is from other TADs (**Methods**); a lower value for DBI indicates a more distinct, better-separated cluster. DCC measures the difference between intra-TAD interaction counts and inter-TAD interaction counts, with higher difference associated with better TADs. For each TAD-finding algorithm, we estimated the proportion of TADs with significantly better DBI or DCC value than randomly shuffled TADs. GRiNCH and HiCseg yield the highest proportion of TADs with significantly better DBI or DCC values compared to randomly shuffled TADs (**Figure 2A**), suggesting these methods provide the most coherent set of TADs.

**Table 1:**
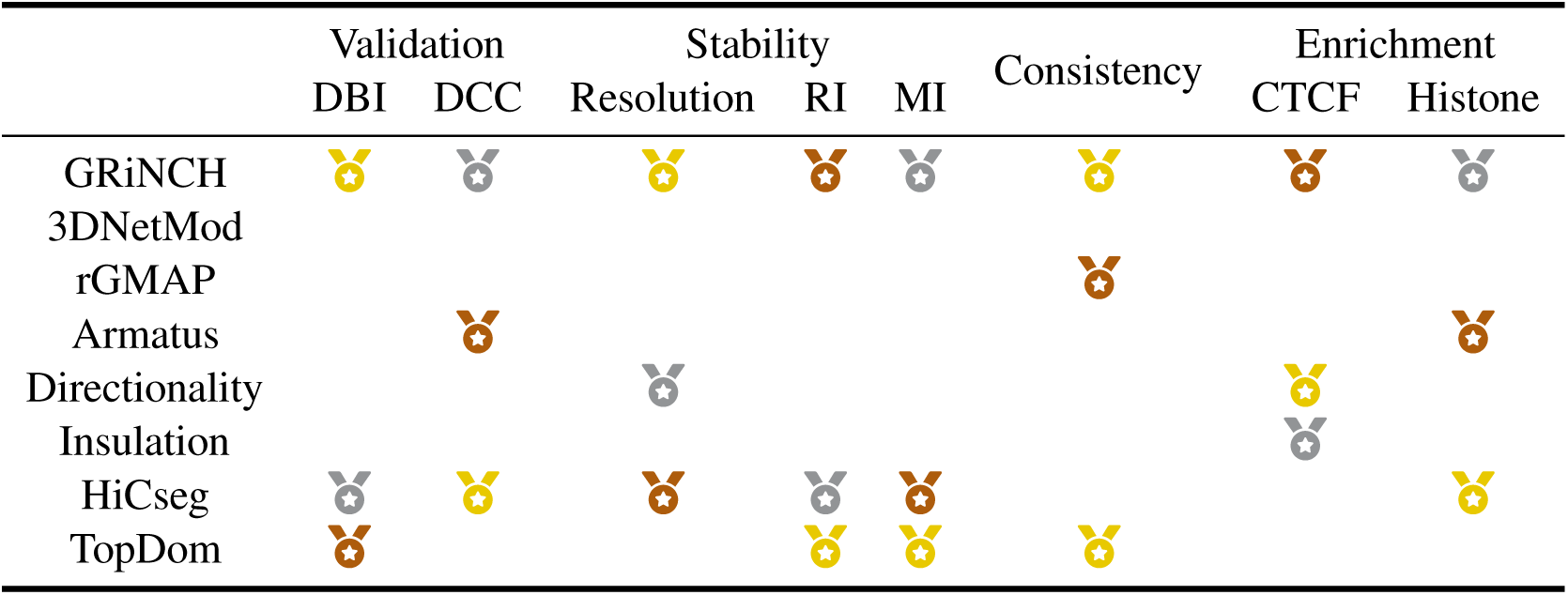
Shown are different criteria of evaluation. A medal denotes whether the given TAD-calling method is among the top 3 methods for a particular criteria (gold/yellow: 1st place; silver/grey: 2nd place; bronze/brown: 3rd place). DBI: proportion of TADs with significant Davies-Bouldin Index; DCC: proportion of TADs with significant Delta Contact Counts; Resolution: stability of median TAD size to Hi-C resolution; RI, MI: stability to depth and sparsity of input data, measured by Rand Index (RI) and Mutual Information (MI); Consistency: a group of methods yielding TADs with highest similarity, with gold for the pair of methods with highest similarity according to hierarchical clustering; CTCF: fold enrichment of CTCF and cohesin elements in TAD boundaries; Histone: proportion of TADs with significant mean histone signal. See **Table S1** and Supplementary Data for more details.

**Figure 2:**
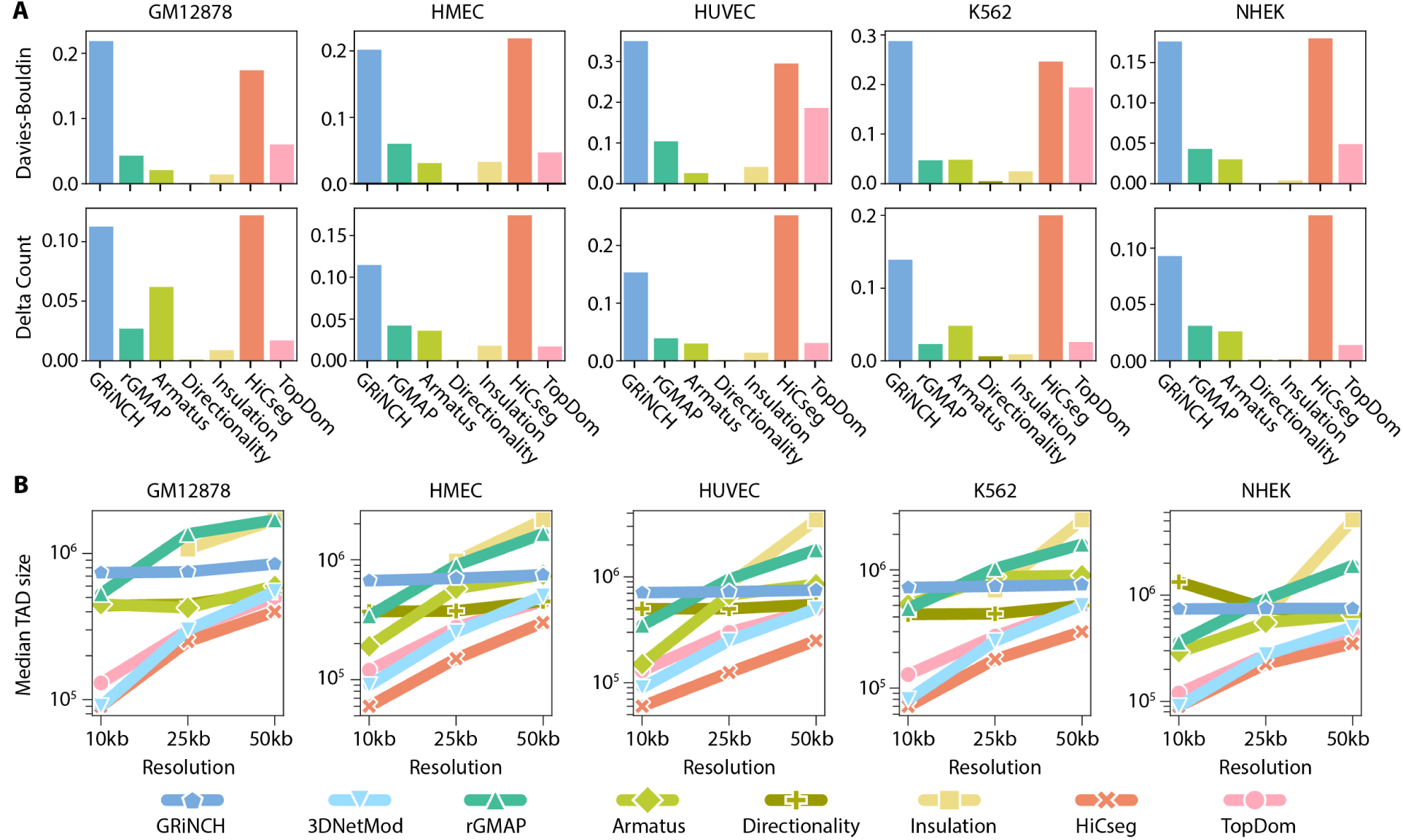
Characterizing TADs with internal validation metrics and median TAD size. **A**. Proportion of TADs with empirical p-value *<* 0.05 for internal validation metrics Davies-Bouldin Index and Delta Count, shown for GRiNCH and six other methods. Note: 3DNetMod outputted overlapping TADs, even when run under non-hierarchical settings and was excluded from this analysis which involves TAD randomization/shuffling. Treating each TAD as a cluster of genomic regions, we evaluate how distinct each cluster is to other clusters (Davies-Bouldin Index) and how much higher the intra-cluster interactions are compared to inter-cluster interactions (Delta Count). The p-value of each cluster’s metric value is derived against the empirical distribution of the metric values in randomly shuffled TADs. The higher the bar, the better a method. **B**. The median size of TADs identified by different methods from different Hi-C resolutions. A method is considered stable to the resolution (size of the region) of the data if the median TAD size does not change substantially with the resolution, given the same user-defined parameter settings.

Many TAD-calling methods are sensitive to the input data resolution (size of genomic region), with the resulting TAD lengths varying greatly as a function of resolution [28]. A robust method is expected to yield consistent length distribution of TADs when given the same user-specified parameter settings, regardless of the change in resolution. Therefore, we next assessed the ability of GRiNCH and the seven TAD calling methods for their ability to recover stable TADs across different resolutions, 10kb, 25kb, and 50kb. When comparing the median length of TADs across different resolutions (**Figure 2B**), GRiNCH and Directionality Index are the most stable, with the exception of NHEK where Directionality index learns longer TADs at 10k resolution. This suggests that GRiNCH is robust to different resolutions, recovering consistently-sized TADs across different resolutions.

TAD-calling methods can be sensitive to the sparsity of the Hi-C matrices due to low sequencing depth [28]. To assess the robustness of each method to low-depth, sparse datasets with many zero entries, we first took the highest-depth dataset (GM12878, 86 million reads total) and downsampled to the depth and sparsity level of lower-depth data from other cell lines (e.g. K562, the second “deepest” cell line with 16 million reads). We then compared the similarity of the TADs from the original high-depth data and those from the downsampled counterpart (**Figure 3A, Methods**). We utilized metrics that can quantify the similarity of pairs of clustering results: Rand Index and Mutual Information (**Methods**). Intuitively, Rand Index is a measure of cluster membership consistency; it measures whether two data points (in our case, two genomic regions) that belonged to the same cluster (TAD) in one clustering result also stayed together in the other result, and whether two data points that belonged to different clusters stayed separate. Rand Index ranges from 0 to 1, with 1 being perfect concordance. Mutual Information is an informational-theoretic metric measuring the dependency between two random variables, where each variable indicates a clustering result. A Mutual Information of 0 indicates complete disagreement and the higher the Mutual Information value the better the agreement between the corresponding clustering results. Based on Rand Index, TopDom, HiCseg, and GRiNCH yield the most reproducible TADs across different depths, particularly at the lower depths of HMEC, HUVEC, and NHEK cell lines. Based on Mutual Information, TopDom is the most consistent followed by GRiNCH and HiCseg. Other methods were generally less consistent based on the Mutual Information metric.

**Figure 3:**
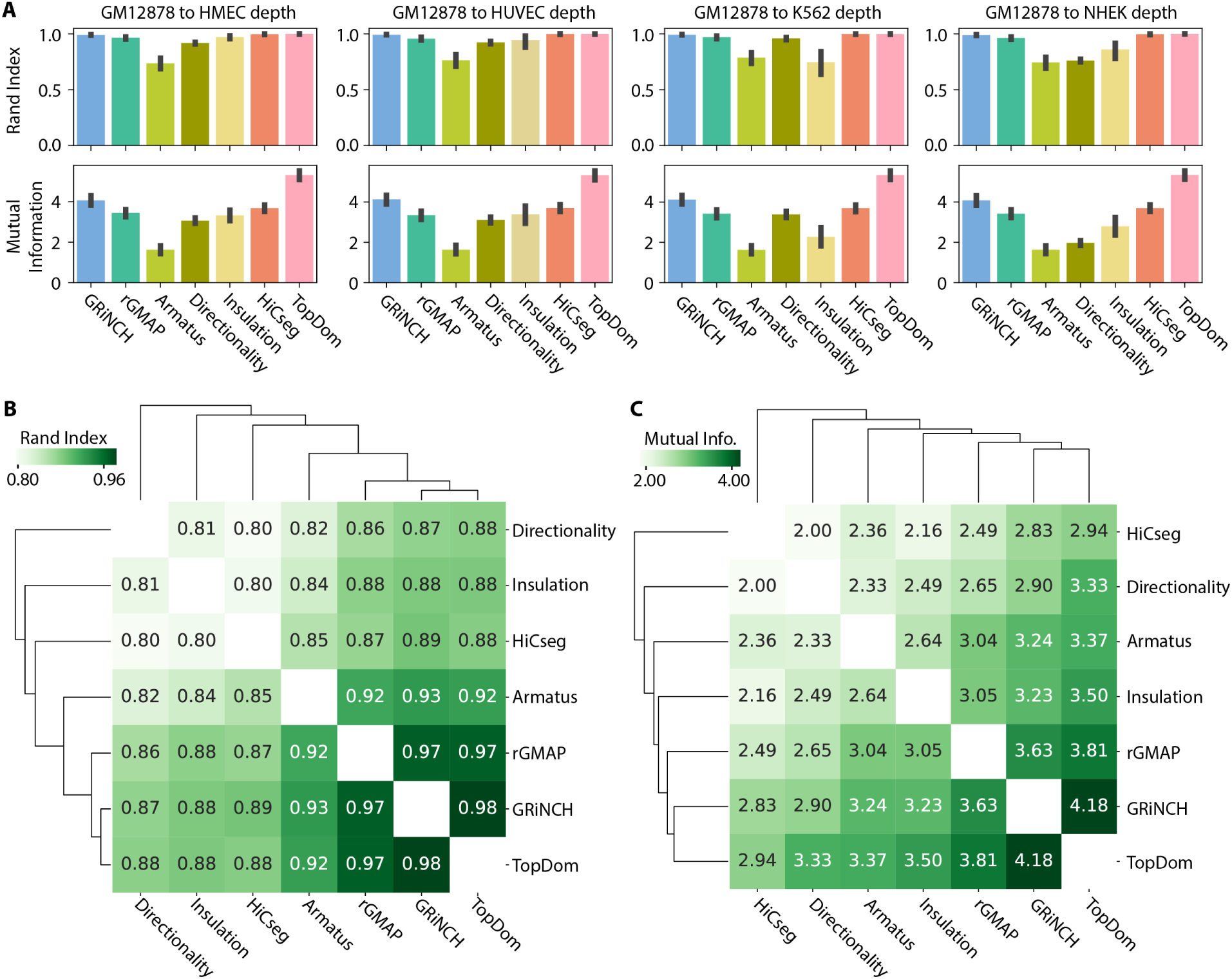
Evaluating the stability and similarity of different TAD-calling methods. **A**. The mean similarity, across chromosomes, between TADs from high-depth GM12878 dataset and TADs from low-depth GM12878 datasets obtained by downsampling the GM12878 dataset to different depths observed in our five cell-line dataset. The similarity of the TADs is measured by Rand Index and Mutual Information. The error bar denotes the standard deviation from the mean. **B**. Similarity of TADs from pairs of TAD-calling methods (e.g. GRiNCH vs. TopDom), measured by Rand index. The higher the number, the higher the similarity. **C**. Similarity of TADs from pairs of TAD-calling methods measured by Mutual information. Note: 3DNetMod outputted overlapping TADs, even when run under non-hierarchical settings; it was excluded from this analysis because of the requirement of distinct within-TAD measurements.

A third hindrance in the interpretation of results from TAD finding methods is the disagreement on the TAD definitions [28, 29]. Hence, we further evaluated whether different TAD-calling methods yielded relatively similar TADs, and which sets of methods yielded the most similar TADs to one another. Here again, we used Rand Index and Mutual Information as metrics to compare the sets of TADs from different methods. All pairwise comparisons of TAD-calling methods yielded high values of Rand Index (*>*0.8) and high Mutual Information (**Figure 3B,C**). Furthermore, GRiNCH and TopDom yield the most similar sets of TADs, followed by rGMAP across all cell lines. This pattern is fairly consistent even when analyzed for each cell line individually (**Figure S2**).

To summarize, our internal validation and stability analysis showed that the top performing methods depends upon the evaluation criteria. However, GRiNCH is among the top performing methods for all the criteria we examined (**Table 1**), producing TADs that are as good or better than existing methods and are stable to varying resolution and depth.

### GRiNCH TADs are enriched in architectural proteins and histone modification signals

We next characterized GRiNCH TADs as well as TADs from other methods for their ability to capture well-known one-dimensional signal enrichment patterns (**Table 1**). In particular, one hallmark of TADs is the enrichment of architectural proteins such as CTCF and cohesin elements (RAD21, SMC3) on the boundaries of TADs [29, 38]. We tested the TAD boundaries from each method for the enrichment of peaks of CTCF, RAD21, and SMC3 in the five Rao et al. cell lines with Hi-C data (**Figure 4A, Methods**). All methods identified boundaries enriched for peaks of these proteins; however, the methods varied in their relative performance across cell lines. GRiNCH TAD boundaries have comparable or better enrichment as the other top performing methods, namely, Directionality Index and Insulation Score in most cell lines, and HiCseg in K562 and Huvec. All these methods including GRiNCH have significantly higher enrichment than 3DNetMod, rGMAP, Armatus across different cell lines.

**Figure 4:**
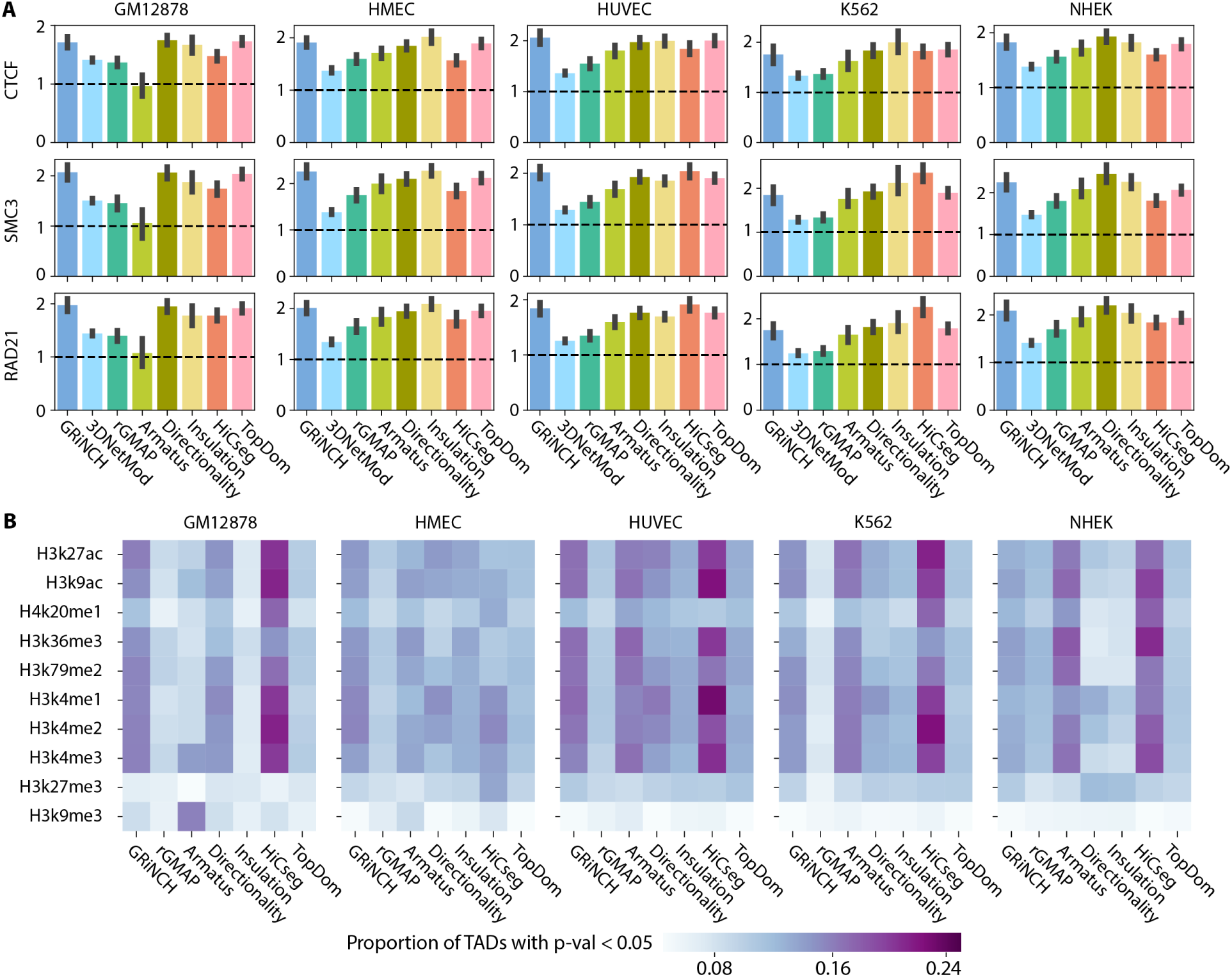
Evaluating the quality of TADs from different TAD-calling methods using enrichment of boundary elements and regulatory signals. **A**. Fold enrichment of binding signals of architectural protein in TAD boundaries. Shown are the mean fold enrichment of CTCF ChIP-seq peaks and accessible motif instances of cohesin proteins, RAD21 and SMC3, estimated across multiple chromosomes. The error bar denotes the standard deviation from the mean. **B**. Proportion of TADs with significant mean histone modification signal (i.e. empirical p-value *<* 0.05). The darker the entry the higher the proportion of TADs with significant histone enrichment. The average ChIP-seq signal for each histone modification mark was taken from within each TAD; the p-value of each TAD is derived from an empirical null distribution of mean signals in randomly shuffled TADs. Note: 3DNetMod outputted overlapping TADs, even when run under non-hierarchical settings; and it was excluded from this analysis as it involves TAD randomization/shuffling.

As histone modifications have been shown to be associated with three-dimensional organization [39], we next measured the proportion of TADs with significant levels of mean histone modification signals (**Figure 4B**) compared to randomly shuffled TADs (**Methods**). The histone modification signals include promoter-(H3K4me3, H3k4me2), elongation-(H3K79me2, H3k36me3), and enhancer-associated marks (H3K27ac), and repressive chromatin marks (H3K27me3). A larger proportion of GRiNCH TADs, along with Armatus and HiCseg TADs, are consistently enriched for the activating histone makes such as H3K27ac, and the elongation marks, H3K36me3 and H3K79me2. Interestingly, with the exception of GM12878, the enrichment of histone marks in the TADs from Insulation and Directionality index was much lower than the other methods suggesting these methods tend to find TADs defined by CTCF and might miss other types of TADs [38]. These enrichment metrics show that when considering existing methods, there is a tradeoff in the ability to recover TADs that are associated with CTCF and TADs that are associated with significant histone modifications. However, GRiNCH ranks among the top methods for both criteria suggesting that GRiNCH TADs capture a diverse types of TADs.

### GRiNCH smoothing of low-depth datasets help recover structure and significant interactions

Our analysis so far compared different TAD finding methods for their ability to recover stable and biologically meaningful topological units. However, most Hi-C datasets are sparse, which can influence the TAD predictions significantly. Smoothing the input Hi-C matrix to impute missing values can enhance the visualization of topological units on the matrix and improve the agreement among biological replicates [30, 31]. Unlike existing TAD-calling methods, the matrix factorization framework of GRiNCH provides a natural matrix completion solution that can generate a smoothed version of the sparse input Hi-C matrix. We next compared GRiNCH’s smoothing functionality to common smoothing techniques such as mean filter and Gaussian filter, which are used in imaging domains and also for Hi-C data [30]. We used two metrics to assess the quality of smoothing: (a) recovery of TADs and (b) recovery of significant interaction after smoothing downsampled data. To perform these comparisons, we again used the downsampled GM12878 datasets.

To assess TAD recovery, we identified TADs on the original high-depth GM12878 dataset and compared them to the TADs identified in the downsampled and smoothed data matrices using Rand Index and Mutual Information. Here, to avoid any bias in our interpretation, we used the Directionality Index method to call TADs. We find that using both Rand Index and Mutual Information, TADs recovered on GRiNCH smoothed matrices are the most similar to the TADs from the high-depth dataset across different parameter settings of the mean filter and Gaussian filters (**Figure 5A**). The usefulness of GRiNCH is more apparent for lower-depth datasets such as NHEK. To assess the recovery of significant interactions, we applied Fit-Hi-C [40] on the original GM12878 dataset and on the downsampled and smoothed datasets to identify significant interactions (q-value *<* 0.05). Treating the significant interactions in the original high-depth dataset as the ground truth, we measured precision and recall as a function of the statistical significance of interactions from the smoothed datasets and computed the Area Under Precision-Recall curve (AUPR). The higher the AUPR, the better the recovery of significant interactions after smoothing. GRiNCH has the highest AUPR compared to mean filter and Gaussian filter (**Figure 5B**) across multiple parameter configurations. Overall, our experiments suggest that GRiNCH offers superior smoothing functionality compared to standard smoothing techniques enabling better recovery of TAD structures and long-range interactions.

**Figure 5:**
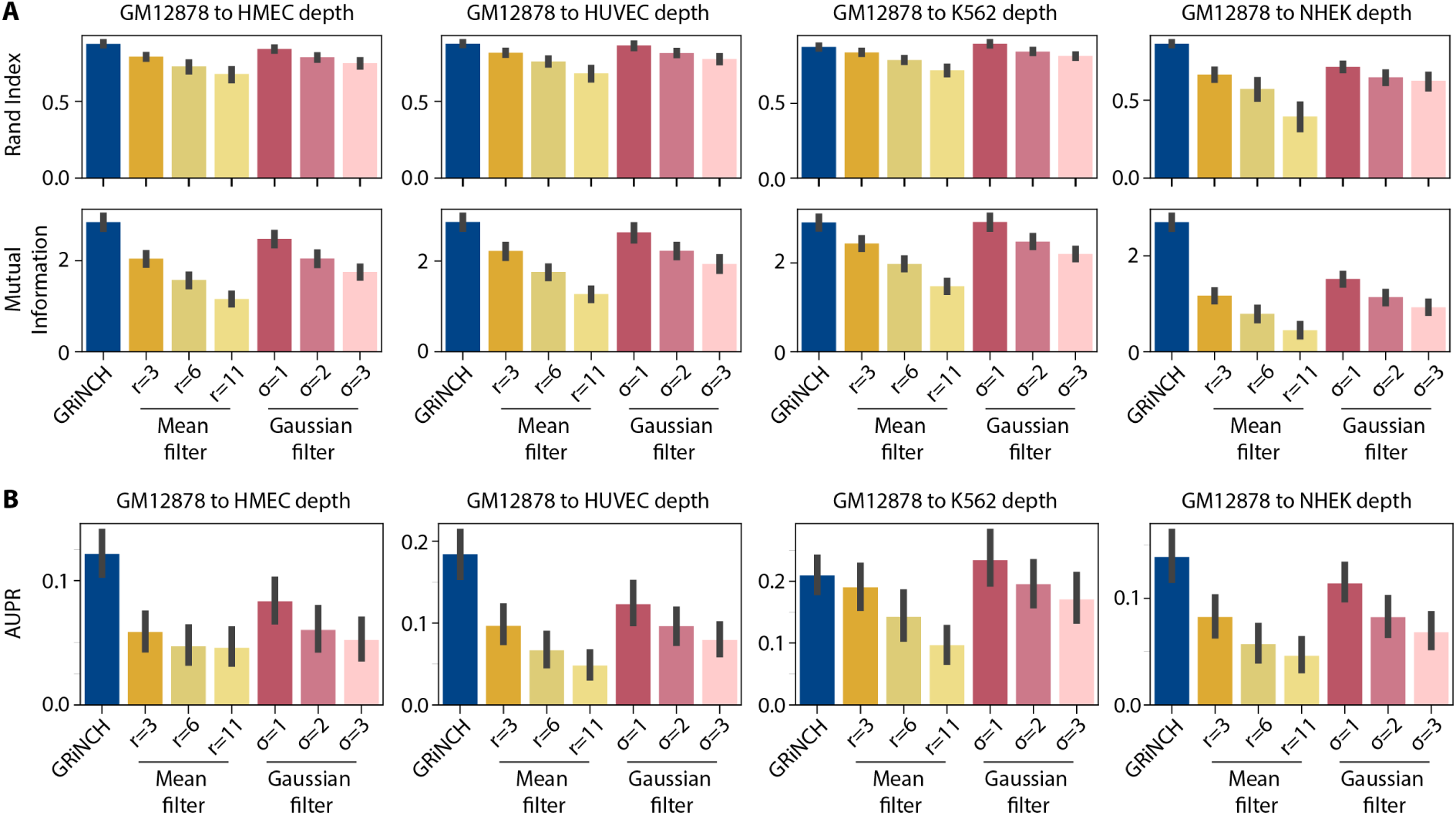
Evaluating the benefits of smoothing in GRiNCH. **A**. Recovery of Directionality Index TADs in downsampled then smoothed data. Shown is the mean similarity (measured by Rand Index and Mutual Information) between Directionality Index TADs from high-depth GM12878 dataset and Directionality TADs from downsampled datasets smoothed by different methods (GRiNCH, Mean Filter, Gaussian Filter). The mean is computed across chromosomes and the error bar denotes deviation from the mean. Directionality was used as a TAD-calling method independent of any of the smoothing methods, i.e., GRiNCH. **B**. Recovery of Fit-Hi-C significant interactions, as measured by the Area Under Precision-Recall curve (AUPR), with precision and recall measured for significant interactions from downsampled and smoothed datasets against the “true” interactions defined as the significant Fit-Hi-C interactions from the high-depth GM12878 dataset.

### GRiNCH application to chromosomal organization during development

To assess the value of GRiNCH in primary cells and to examine dynamics in chromosomal organization, we applied GRiNCH to two time-course Hi-C datasets profiling 3D genome organization during (a) mouse neural development [41] and (b) pluripotency reprogramming in mouse [42]. Bonev et al. [41] used high-resolution Hi-C experiments to measure 3D genome organization during neuronal differentiation from the embryonic stem cell state (mESC) to neural progenitor cells (NPCs) and cortical neurons (CNs). We applied GRiNCH on all chromosomes for all three cell types and compared them based on the overall similarity of TADs between the cell lines. Based on the two metrics of Mutual Information and Rand Index, the overall TAD similarity captured the temporal ordering of the cells, with CNs being closer to NPCs and ESCs the most distinct (**Figure S3A**). We next focused on a specific 4Mbp region around the Zfp608 gene, which was found by Bonev et al. as a neural-specific gene associated with a changing TAD boundary. In both NPCs and CNs, GRiNCH predicts a TAD near the Zfp608 gene, which is not present in the mESC state. Zfp608 was also associated with increased expression, and activating marks, H3K27ac and H3K4me3 at these time points, which is consistent with Zfp608 being a neural-specific gene (**Figure 6A**).

**Figure 6:**
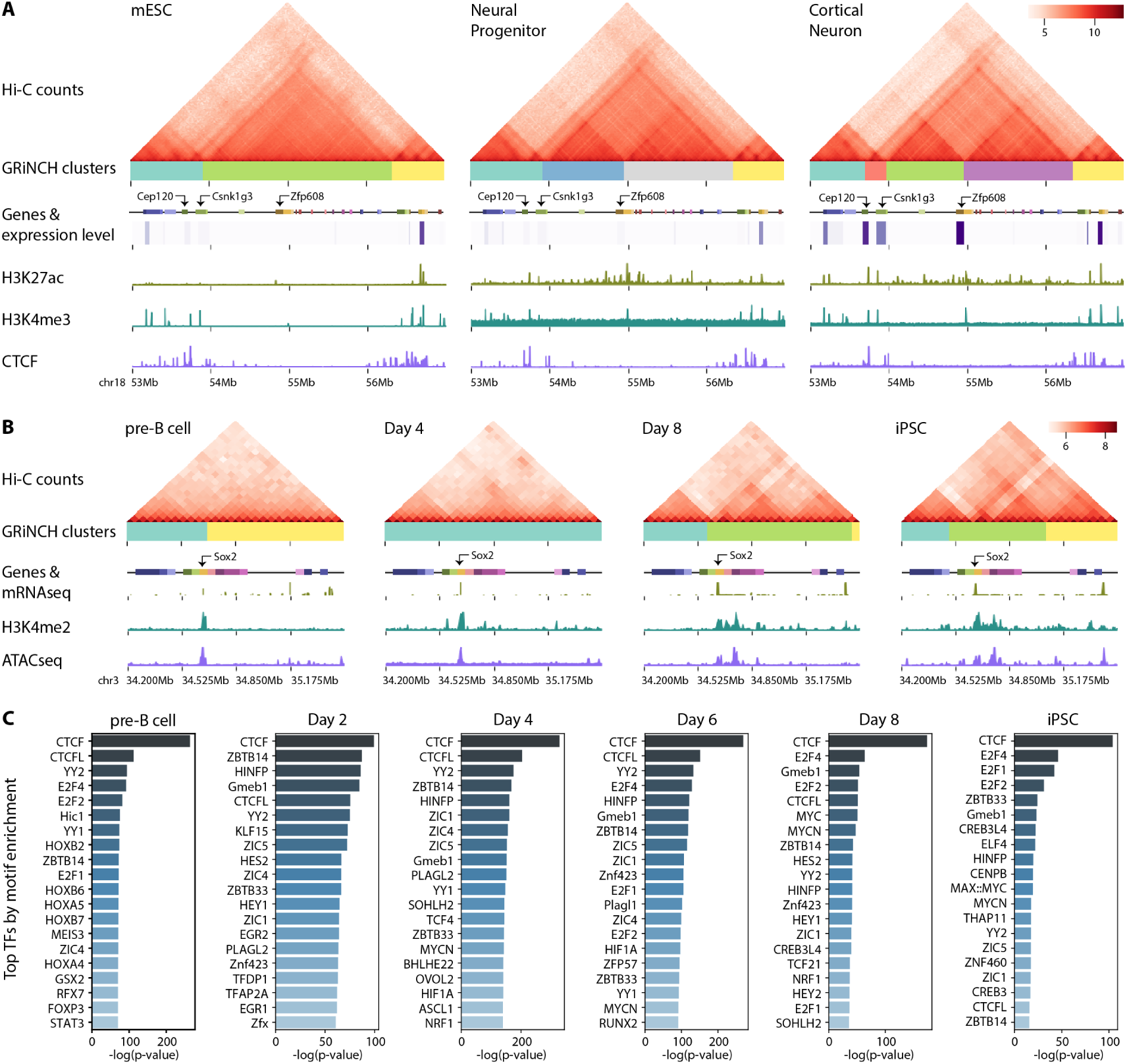
GRiNCH applied to Hi-C datasets along developmental time courses. **A**. Interaction profile near the Zfp608 gene in mouse embryonic stem cells (mESC), neural progenitors (NPC), and differentiated cortinal neurons (CN). Heatmaps are of Hi-C matrices after log2-transformation of interaction counts for better visualization. GRiNCH clusters are visualized as blocks of different colors under the heatmap of interaction counts. Genes in the nearby regions are marked by small boxes, and a heatmap of their corresponding RNA-seq levels (in TPM) is shown underneath each gene. ChIP-seq signals from H3K27ac, H3K4me3, and CTCF are shown as separate tracks. **B**. Interaction profile near the Sox2 gene in mouse pre-B cells, in day 4 and day 8 of reprogramming, and in induced pluripotent stem cell (iPSC). Heatmaps are of Hi-C matrices after log2-transformation of interaction counts for better visualization. GRiNCH clusters are visualized as blocks of different colors under the heatmap of interaction counts. Genes in the nearby regions are marked by small boxes, and peaks of their corresponding RNA-seq levels are shown underneath each gene. ChIP-seq signals from H3K4me2 and ATAC-seq signals are shown as separate tracks. **C**. Top 20 TFs from a collection of 746 TFs ranked based on their motif enrichment in GRiNCH TAD boundaries from the mouse reprogramming time course data. The significance of their fold enrichment was calculated with the hypergeometric test and TFs were ranked by descending negative log p-value.

We next examined another time-course dataset which studied the 3D genome organization during reprogramming of mouse pre-B cells to pluripotent stem cells (PSC), with four intermediate time points (Day 2, 4, 6, and 8, see **Methods**). As in the neural developmental time course, we applied GRiNCH to all chromosomes from each time point and compared the overall 3D genome configuration over time. Here too we observed that time points closer to each other generally had greater similarity in their TAD structure, as well as two different replicates within the same time point displaying even greater TAD similarity (**Figure S4B**). We examined the interaction profile in the 1.3 Mbp around the Sox2 gene, a known pluripotency gene (**Figure 6B**). We see a gradual formation of a boundary around Sox2, which is also associated with concordant increase in expression, accessibility and the presence of H3K4me2, an active promoter mark.

As chromatin accessibility data was also measured at each timepoint during reprogramming, we asked if we could identify additional regulatory proteins that could play a role in establishing TADs (**Methods**). Briefly, we tested the GRiNCH TAD boundaries from each mouse cell type, from pre-B cell to pluripotent cells, for enrichment of accessible motif instances of 746 transcription factors in the JASPAR 2020 core vertebrate motif database [43]. We ranked the TFs based on their significant enrichment in each cell type (**Figure 6C, Table S2**). The top-ranking TF across the cell types was CTCF, which is consistent with its role as an architectural protein in establishing TADs (**Figure 6C**). We also found other factors in the same zinc finger protein family as CTCF [44], such as ZBTB14, Plagl2/1, ZIC1/3/4/5, CTCFL, YY1/2 that were enriched across the cell types. YY1 and YY2 (which are 65 and 56% identical in their DNA and protein sequence respectively in humans [45]), are of interest, as YY1 has been identified as an enforcer of long-range enhancer-promoter loops [46]. Interestingly, we found several hematopoietic lineage factors, such as STAT3 and FOXP3, ranked highly in the pre-B cell TADs compared to other time points. STAT3 is needed for B cell development [47]. FOXP3 is a master regulator of T cells [48], but could be involved in the suppression of B cells. We also found a number of HOX transcription factors, HOXA4, HOXA5, HOXB2, HOXB5, HOXB7, and the transcription factor MEIS3 to be ranked highly in the B cells. The HOX genes depend upon MEIS3 [49] to bind to their targets, supporting the simultaneous enrichment of these factors.

We repeated this analysis for the Rao et al. cell lines (**Table S3**). Here too we found CTCF and YY1/2 proteins highly enriched across cell lines. However, there was lesser degree of cell-line specificity for this dataset. Taken together, this analysis suggests that GRiNCH captures high-quality TADs, which can be used to define global similarities and difference between cell types. Furthermore, the GRiNCH boundary enrichment analysis identified novel transcription factors that could be followed up with downstream functional studies to examine their role in 3D genome organization.

### GRiNCH can be used for a variety of 3D conformation capture technologies

Although Hi-C is still the most widely used technology to map 3D genome structure, recently several new methods have been developed to measure chromosomal contacts on a genome-wide scale [6]. To assess the applicability of GRiNCH to these technologies, we considered two complementary techniques to measure 3D genome organization: Split-Pool Recognition of Interactions by Tag Extension (SPRITE) [9] and HiChIP [37]. SPRITE measures multi-way chromatin interactions, and captures interactions across larger spatial distances than Hi-C. In HiChIP, long-range chromatin contacts are first established *in situ* in the nucleus before lysis; then chromatin immunoprecipitation (ChIP) is performed with respect to a specific protein or histone mark, directly capturing interactions associated with a protein or histone mark of interest [37]. A common property of both technologies is that they generate a contact count matrix, which is suitable for GRiNCH.

We applied GRiNCH to GM12878 contact matrices measured with SPRITE [9], cohesin HiChIP [37], and H3k27ac HiChIP [50]. A visual comparison between these datasets for an 8Mb region of chr8 shows regions of good concordance between datasets (**Figure 7A-D**). We quantified the global similarity of GRiNCH TADs from the four different datasets, for all chromosomes, with Rand Index (**Figure 7**E) and Mutual Information (**Figure 7**F). Interestingly, the GRiNCH TADs from Hi-C are the most similar to those from cohesin HiChIP and this similarity measure is higher than between the two HiChIP datasets. This is consistent with cohesin being a major determinant for the formation of loops detected in HiC datasets. The H3K27ac HiChIP data is as close to Hi-C as it is to cohesin HiChIP. Finally the most distinct set of TADs are identified by SPRITE, which is consistent with SPRITE capturing multi-way interactions and longer-distance interactions. Despite the differences in the cluster, overall the datasets look similar across different platforms (Rand Index *>*0.97). Taken together, this shows that GRiNCH is broadly apply to different experimental platforms for measuring genome-wide chromosome conformation.

**Figure 7:**
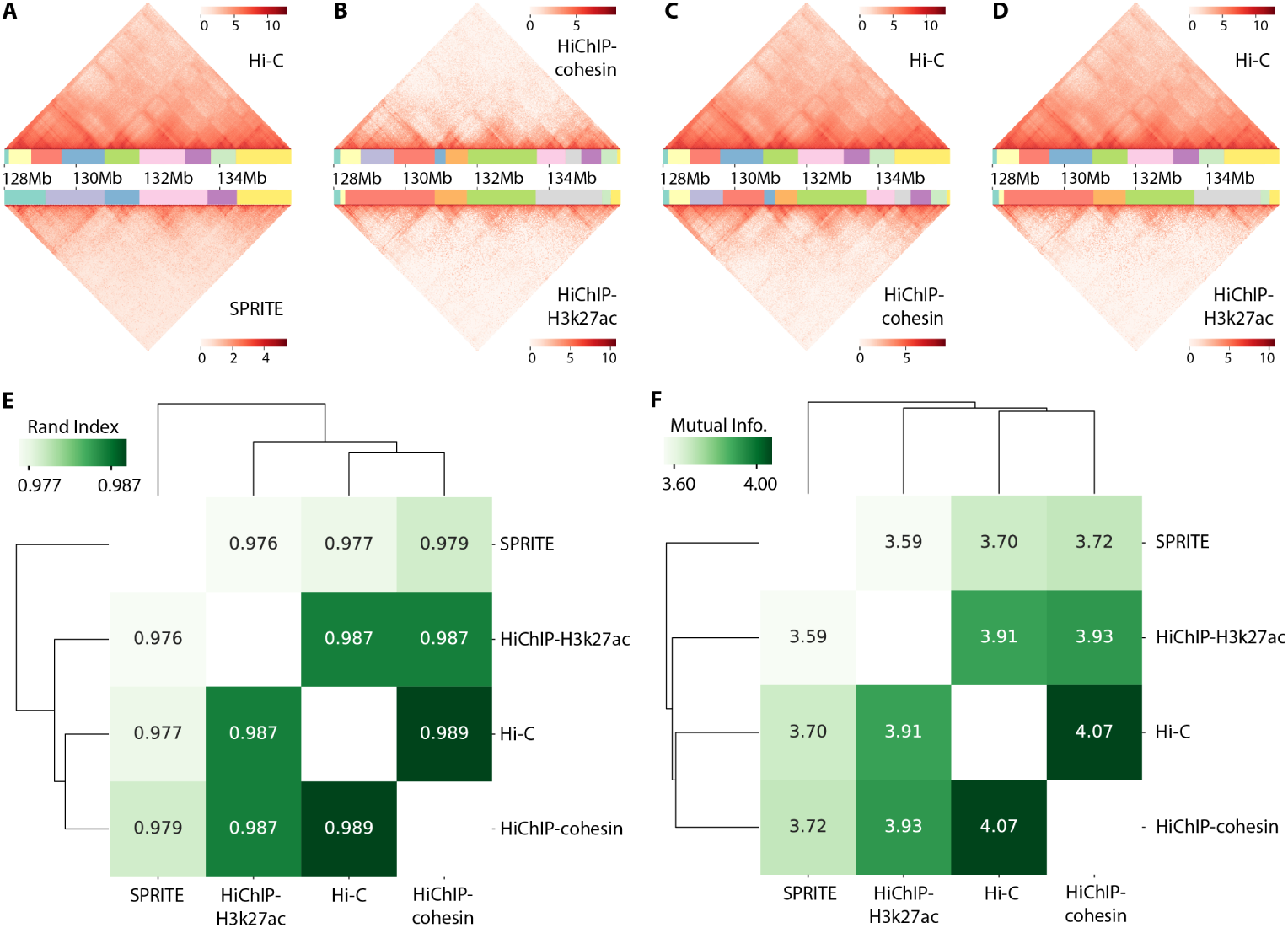
Applying GRiNCH to datasets from different 3D genome conformation capture technologies. Visual comparison of the interaction profile and GRiNCH TADs from a 8Mb region in chr8, GM12878 cell line. GRiNCH TADs are visualized as blocks of different colors under the heatmap of interaction counts. **A**. Hi-C vs SPRITE. The top heatmap and clusters are from Hi-C; bottom from SPRITE. **B**. HiChIP with cohesin (top) vs HiChIP with H3k27ac (bottom). **C**. Hi-C (top) vs HiChIP with cohesin (bottom). **D**. Hi-C (top) vs HiChIP with H3K37ac (bottom). For visualization purposes all interaction counts were log2-transformed. **E**. Measuring the similarity of GRiNCH TADs from Hi-C and other 3D genome conformation capture platform (e.g. SPRITE, HiChIP with cohesin, or HiChIP with H3k27ac) in the same GM12878 cell line, with Rand Index. The dendrogram depicts the relative similarity between samples. **F**. Mutual-Information-based similarity of GRiNCH TADs from Hi-C and other technologies.

## Discussion

We present GRiNCH, a graph-regularized matrix factorization framework that enables reliable identification of high-quality genome organizational units, such as TADs, from high-throughput chromosome conformation capture datasets. GRiNCH is based on a novel constrained matrix factorization and clustering approach that enables recovery of contiguous blocks of genomic regions sharing similar interaction patterns as well as smoothing sparse input datasets.

A lack of gold standards for TADs emphasizes the need to probe both the statistical and biological nature of inferred TADs. Through extensive comparison of GRiNCH to existing methods with good performance in other benchmarking studies, we identified strengths and weaknesses of existing approaches. In particular, methods like Insulation Score identify TADs that are generally more enriched for signals such as CTCF and cohesin; however, when comparing statistical properties such as stability across resolutions and cluster coherence, this method does not necessarily rank the best. GRiNCH was among the top methods for both criteria, identifying clusters of genomic regions with high degree of similarity in their interaction profiles, stable to low-depth, sparse datasets, and enriched in architectural proteins and histone modification signals with known roles in chromatin organization.

A unique advantage of GRiNCH lies in its smoothing capability via matrix completion. Smoothing has been an independent task from TAD-calling and a key processing step in downstream analysis of Hi-C data (e.g. measuring reproducibility or concordance between Hi-C replicates [31]). We find that GRiNCH smoothing outperforms existing smoothing methods (mean filter and Gaussian filter) in its ability to retain TAD-level and interaction-level features of the input Hi-C data. Furthermore, GRiNCH is applicable to datasets from a wide variety of platforms, including SPRITE and HiChIP. Application of GRiNCH shows that Hi-C and HiChIP datasets capture more similar topological units than SPRITE. Interestingly, TADs from Hi-C and cohesin HiChIP are much closer than the two HiChIP datasets we compared. This shows that GRiNCH is capturing TADs that are reproducible across platforms. To study the ability of GRiNCH to identify dynamic topological changes along a time course, we applied GRiNCH to published developmental time-course datasets. GRiNCH recapitulated global temporal relationships in 3D organization and also transitions in topological units around key developmental genes. Thus, GRiNCH should be broadly applicable for analysis of chromosome conformation capture datasets with different experimental design, sequencing depths, and platforms.

The 3D organization of the genome is determined through a complex interplay of architectural proteins such as CTCF, cohesin elements, and other transcription factors such as WAPL [51]. Application of GRiNCH to Hi-C datasets representing cell lines and temporally related conditions identified known and novel transcription factors that could be important for establishing these boundaries in a cell-type-specific or generic manner. In particular, we recovered YY1/2 proteins that have been shown to interact with CTCF to establish long-range regulatory programs during lineage commitment [52]. Among the novel factors that were present in both the cell lines as well as the mouse reprogramming dataset, were several zinc finger proteins, e.g. PLAGL1, ZIC1, ZIC4/5, ZBTB14; such proteins can be investigated for their role in establishing organizational units in mammalian genomes. We also found several factors that were specific to cell lines and time points. For example, FOXI1, a forkhead protein, was ranked highly in K562. Forkhead proteins are involved in genome organization and replication timing in yeast [53] and zebra fish [54], but their role in mammalian genome organization is not well known. The time course data identified additional unique TFs that are likely involved in determining specific lineages, e.g. STAT3, MEIS3, FOXP3 and HOX genes in pre-B cells. HOX genes [55], FOXP3 [56], and STAT3 [47] in particular have been shown to play critical roles in B cell and T cell development. While MEIS1 and MEIS2 are involved in the hematopoietic lineage, MEIS3 specifically is involved in the binding of HOX TFs to target genes in the brain [49]. Therefore the simultaneous enrichment of MEIS3 and HOX sites is consistent with HOX proteins requiring MEIS3 for binding; however, its specific role in the hematopoietic lineage is yet unknown. Investigating the interactions of these proteins with well-known architectural proteins such as CTCF and cohesin could provide mechanistic insight into the factors governing 3D genome organization [29, 57].

There are several directions of future work that are natural extensions to our framework. Although our current approach of analyzing temporal organization in time-course data extracted interesting biological insights, TADs are identified independently for each time point, making it difficult to study the conservation and specificity of individual TADs. One area of future work is to allow joint identification of TADs or similar structural units across multiple conditions [58, 59]. Another direction is to leverage one-dimensional features to potentially inform the TAD-finding algorithm. The GRiNCH framework makes use of a distance dependence graph of regions; however, one could use the similarity of epigenomic profiles to construct an additional graph to constrain the NMF solution.

In conclusion, GRiNCH offers a unified solution, applicable to diverse platforms, to discover reliable and biologically meaningful topological units, while handling sparse high-throughput chromosome conformation capture datasets. The outputs from GRiNCH can be used to predict novel boundary elements, enabling us to test possible hypotheses of other mechanisms for TAD boundary formation. We have made GRiNCH available at roy-lab.github.io/grinch, with a comprehensive installation and usage manual. As efforts to map the three-dimensional genome organization expand to more conditions, platforms, and species, a method such as GRiNCH will serve as a powerful analytical tool for understanding the role of genome 3D organization in diverse complex processes.

## Materials and Methods

### Graph-regularized Non-negative Matrix Factorization (NMF) and Clustering for Hi-C data (GRiNCH) framework

GRiNCH is based on a regularized version of non-negative matrix factorization (NMF) [35] that is applicable to high-dimensional chromosome conformation capture data such as Hi-C (**Figure 1**). Below we describe the components of GRiNCH: NMF, graph regularization, and clustering for TAD identification.

### Non-negative matrix factorization (NMF) and graph regularization

Non-negative matrix factorization is a popular dimensionality reduction method that aims to decompose a non-negative matrix, X ∈ ℝ^(*n*×*m*)^ into two lower dimensional non-negative matrices, U ∈ ℝ^(*n*×*k*)^ and V ∈ ℝ(*n*×*k*), such that the product X^*^ = UV^T^, well approximates the original X. We refer to the U and V matrices as factors. Here *k << n, m* is the rank of the factors and is user-specified.

In application of NMF to Hi-C data, we represent the Hi-C data for each chromosome as a symmetric matrix X = [*x*_*ij*_] ∈ ℝ^(*n*×*n*)^ where *x*_*ij*_ represents the contact count between region *i* and region *j*. We note that in the case of a symmetric matrix, U and V are the same or related by a scaling constant.

The goal of NMF is to minimize the following objective: ‖X − UV^*T*^ ‖^2^, s.t. U ≥ 0, V ≥ 0 [32]. A number of algorithms to optimize this objective have been proposed; here we used the multiplicative update algorithm, where the entries of U and V are updated in an alternating manner each iteration:

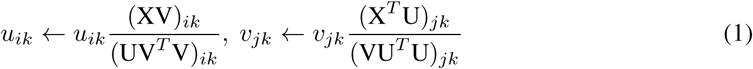

Here *u*_*ik*_ corresponds to the *i*^th^ row of column U(:, *k*) and *v*_*jk*_ corresponds to the *j*^th^ row of column V(:, *k*).

Standard application of NMF to Hi-C data is ignorant of the strong distance dependence of the count matrix, that is, genomic regions that are close to each other tend to interact more with each other. To address this issue we apply an constrained version of NMF with graph regularization, where the graph represents additional constraints on the row (and/or column) entities [35]. Graph regularization enables the learned columns of U and V to be smooth over the input graph. In our application of NMF to Hi-C data, we define a graph composed of genomic regions as nodes, with edges connecting neighboring regions in the linear chromosome, where the size of the neighborhood is an input parameter. Specifically, we define a symmetric nearest-neighbor graph, W:

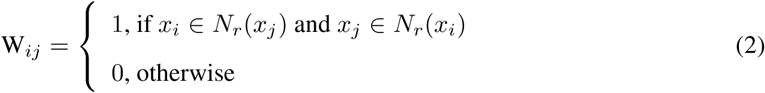

where *N*_*r*_(*x*_*i*_) denotes *r* nearest neighbors in linear distance to region *x*_*i*_. Graph regularized NMF has the following objective:

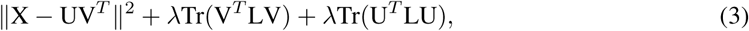

where D is a diagonal matrix whose entries are column (or row, since W is symmetric) sums of W, i.e., D_*ii*_ = Σ_*j*_ W_*ij*_. L = D − W denotes the graph Laplacian and encodes the graph topology. The second and third terms are the regularization term and measures the smoothness of *U* and *V* with respect to the graph. Here *λ* is the regularization hyperparameter. This new objective has the effect of encouraging the factors to be smooth on the local neighborhood defined by the graph. Accordingly, the multiplicative update rule from (1) gains regularization terms [35]:

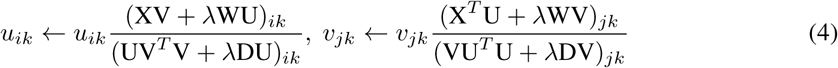

Both *r* (neighborhood radius) and *λ* are parameters that can be specified, with *λ* setting the strength of regularization (*λ* = 0 makes this equivalent to basic NMF). See section on “Estimating GRiNCH hyper-parameters” below.

### Chain-constrained *k*-medoids clustering for clustering assignment and TAD calling

The factors U (or V) can be used to extract clusters of the row (or column) entities of the input matrix. Because X is symmetric in our application, either U or V can be used to define the clusters (the factors are equivalent up to a scaling constant). Assuming we use U, there are two common approaches for finding clusters from NMF factors: (1) assign each row entity *i* to its most dominant factor, i.e., assign it to cluster *c*_*i*_ = argmax_*j*∈{1,…,*k*}_*u*_*ij*_, or (2) apply *k*-means clustering on the rows of U. However, both approaches fall short in our application. The first approach is sensitive to extreme values which can still be present in the smoother factors, yielding non-informative clusters. Furthermore, neither approach reinforces contiguity of genomic regions in each cluster along their chromosomal position. As a result, a single cluster could potentially contain genomic regions from two opposite ends of the chromosomes instead of being a contiguous local structural unit. To address this problem, we apply chain-constrained *k*-medoids clustering. *k*-medoids clustering is similar to *k*-means clustering, except that the “center” of each cluster is always an actual data point, rather than the mean of the datapoints in the cluster. In its chain-constrained version (Algorithm 1), adopted from spatially connected *k*-medoids clustering [60]: each cluster grows outwards from initial medoids along the linear chromosomal coordinates. The algorithm assigns a genomic region to a valid medoid region either upstream or downstream along the chromosome, ensuring the contiguity of the clusters and resilience to noise or extreme outliers provided by using a robust ‘median’-like cluster center rather than a ‘mean’-like center used in *k*-means clustering.

#### Algorithm 1: Chain-constrained *k*-medoids clustering

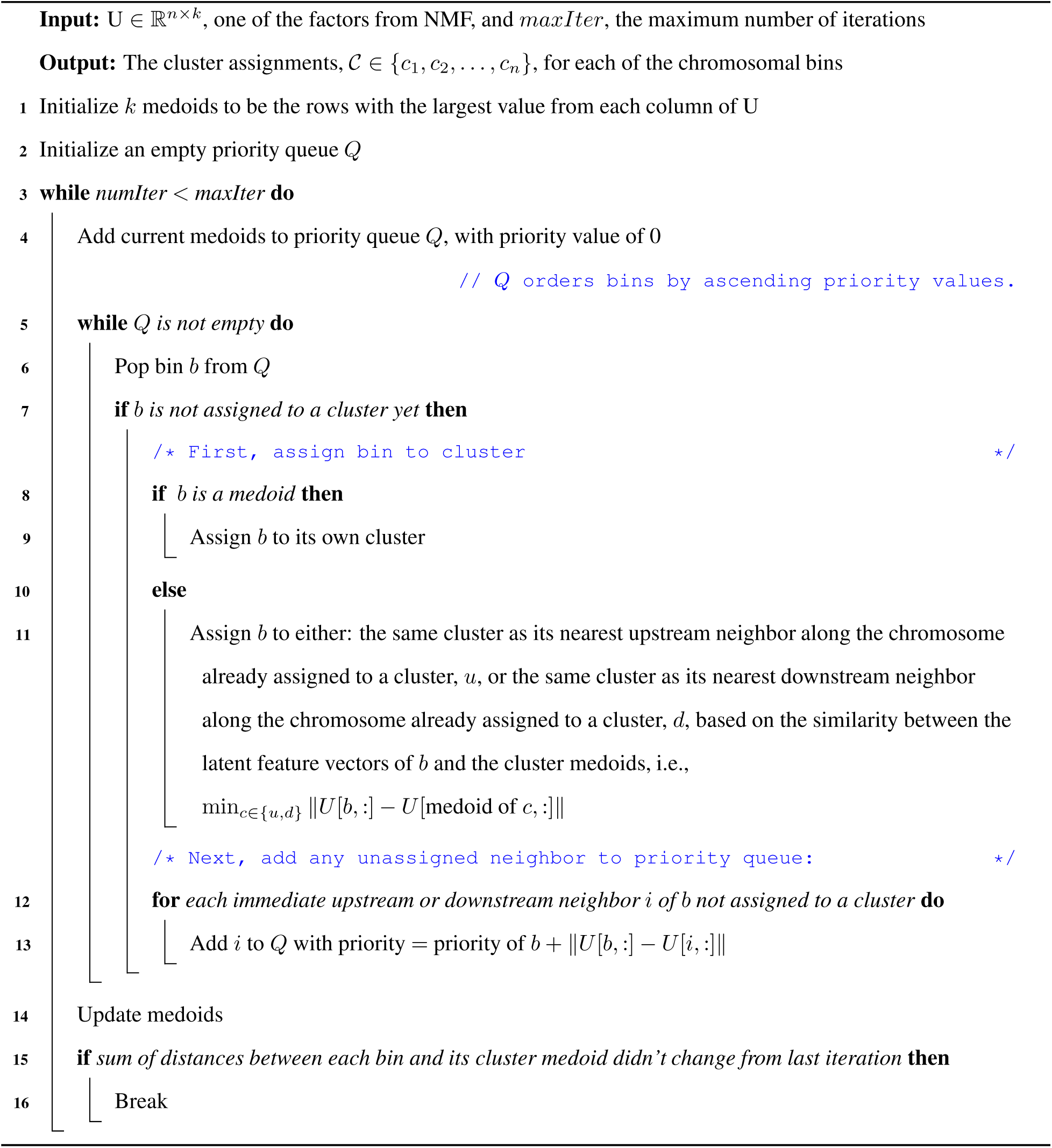

### Selecting GRiNCH hyperparameters

GRiNCH has three hyper-parameters: (a) *k*, the dimension of the lower-dimensional factors which can alternately be viewed as the number of latent features or clusters, (b) *r*, the radius of the neighborhood in the graph used for regularization, and (c) *λ* controlling the strength of regularization.

The parameter *k* determines the number of latent features to recover and the resulting number of GRiNCH TADs. We can yield subTAD-, TAD-, or metaTAD-scale clusters (**Figure S5**A) by setting *k* such that the expected size of a cluster is 500kb, 1Mb, or 2Mb, i.e., *k* equals the given chromosome’s length divided by the expected size. We find that a larger portion of subTAD-scale clusters (i.e. expected TAD size = 500kb) have significant internal validation metric values (**Figure S5**B). SubTAD-scale clusters tend to be more stable to depth and sparsity (**Figure S5**C), and are also more enriched in boundary elements like CTCF (**Figure S6**A). As a tradeoff, higher proportion of metaTAD-scale clusters (i.e. expected cluster size = 2Mb) are enriched in histone modification marks (**Figure S6**B). Based on the use case of GRiNCH, *k* can be set dynamically by the user; by default, GRiNCH sets *k* such that the expected size of a cluster is 1Mb, or at TAD-scale.

For regularization strength, *λ* ∈ {0, 1, 10, 100, 100} were considered, with *λ* = 0 equivalent to standard NMF without regularization. For neighborhood radius, *r* ∈ {25K, 50K, 100K, 250K, 500K,1M} were considered, where *r* = 100K in a Hi-C dataset of 25Kb resolution will use 4 bins on either side of a given region as its neighbors. We find that some regularization, with *λ* = 1, yields better CTCF enrichment than other *λ* values **(Figure S1**A). With regularization, a neighborhood radius of 100Kb or larger yields higher CTCF enrichment (**Figure S1**B). Based on these results, the default regularization parameters for GRiNCH are set at *λ* = 1 and *r* =250kb.

### Stability and initialization of NMF

The NMF algorithm is commonly initialized with random non-negative values for the entries of U and V. The initial values can significantly impact the final values of U and V [61]. This leads to instability of the final factors hinging on the randomization schemes or changing seeds. To address the instability, we used Non-Negative Double Singular Value Decomposition (NNDSVD), which initializes U and V with a sparse SVD approximation of the input matrix X [62]. Since the derivation of exact singular values can considerably slow down the initialization step, we use a randomized SVD algorithm which derives approximate singular vectors [63]. NNDSVD initialization with randomized SVD results in lower loss, i.e. factors that can better approximate the original Hi-C matrix, in fewer iterations (**Figure S7**A,B), and more stable results than direct random initialization (**Figure S7**C,D).

## Datasets used in experiments and analysis

### High-throughput chromosome conformation capture data

We applied GRiNCH to SQRTVC-normalized Hi-C matrices from five cell lines, GM12878, NHEK, HMEC, HUVEC, and K562 at 10kb, 25kb, and 50kb resolution from Rao et al. [36] (GEO accession: GSE63525). We also applied GRiNCH to datasets from other technologies that capture the 3D genome structure and chromatin interactions: Split-Pool Recognition of Interactions by Tag Extension (SPRITE) [9] and HiChIP [37]. We used the SPRITE data for GM12878 cell line (GEO accession: GSE114242). For HiChIP, we applied GRiNCH to the contact matrices from HiChIP with cohesin (GEO accession: GSE80820) [37] and HiChIP with H3k27ac (GEO accession: GSE101498) [50].

We applied GRiNCH to two different mouse developmental time course data: (a) neural differentiation Hi-C data from embryonic stem cells (mESC), neural progenitors (NPC), and cortical neurons (CN) [41] and (b) Hi-C data from reprogramming pre-B cells to induced pluripotent state [42] (GEO accession: GSE96553). For (a) neural differentiation dataset, Juicer Straw tool [64] was used to obtain 25kb Hi-C matrices with vanilla-coverage square-root normalization (original GEO accession: GSE96107). For (b) pluripotency reprogramming, we applied GRiNCH to published normalized Hi-C data from pre-B cells, B*α* cells, day 2 of reprogramming, day 4, day 6, day 8, and finally, pluripotent cells.

### ChIP-seq, DNaseq, ATACseq, and motif datasets

To interpret the GRiNCH results and for comparison to other methods, we obtained a number of ChIP-seq datasets. For CTCF, ChIP-seq narrow-peak datasets available as ENCODE Uniform TFBS composite track [65] were downloaded from the UCSC genome browser (wgEncodeEH000029, wgEncodeEH000075, wgEncodeEH000054, wgEncodeEH000042, wgEncodeEH000063).

As ChIP-seq data for SMC3 and RAD21 is not available in the five cell lines from Rao et al [36], we generated a list of cell-line specific accessible motif sites. Accessible motif sites are defined as the intersection of motif-match regions and DNase-accessible regions in the given cell line. The SMC3 and RAD21 motif matches to the human genome (hg19) was obtained from [66]. To create a union of DNase hotspot regions from replicates within a cell line, BEDtools [67] merge program was used. Finally, the intersection of DNase hotspot regions and motif match regions was calculated for each cell line using BEDtools intersect program. DNase hotspot data was obtained from the ENCODE consortium [68, 69]: ENCFF856MFN, ENCFF235KUD, ENCFF491BOT, ENCFF946QPV, ENCFF968KGT, ENCFF541JWD, ENCFF978UNU, ENCFF297CKS, ENCFF569UYX.

We obtained ChIP-seq datasets for histone modification marks from the ENCODE consortium [68, 69]. To generate genome-wide histone modification levels for each mark, fastq reads were aligned to the human genome (hg19) with bowtie2 [70], and aggregated into a base-pair signal coverage profile using SAMtools [71], and BEDtools [67]. The base-pair signal coverage was averaged within each 25kb bin to match the resolution of Hi-C dataset. The aggregated signal was normalized by sequencing depth within each replicate; the replicates were collapsed into a single value by taking the median.

In order to identify additional novel transcription factors that could play a role in 3D genome organization, we obtained motifs of 746 different transcription factors from JASPAR core vertebrate collection [43]. Next, we obtained obtained their accessible motif match sites to hg19 and mm10 for the five cell lines from [36] using the same process that was used for SMC3 and RAD21 motifs. To identify the accessible motif sites for mouse cells during pluripotency reprogramming [42], we aligned ATACseq fastq reads to the mouse genome (mm10) with bowtie2 [70] and deduplicated with SAMtools [71]. Accessible peaks were called with MACS2 [72]. The ATACseq peaks were then used in place of DNaseq hotspots to find the accessible motif sites as was done for SMC3 and RAD21 motifs.

### TAD calling methods

GRiNCH was benchmarked against 7 other TAD-calling methods: Directionality Index method [23], Armatus [20], Insulation Score method [25], rGMAP [24], 3DNetMod [22], HiCseg [73] and TopDom [74]. For all methods, default or recommended parameters values were used when available. Execution scripts containing the parameter values used for these methods are available to download.

### Directionality index

Directionality index uses a hidden Markov model (HMM) on estimated Directionality Index (DI) scores. The DI score for a genomic region is determined by whether the region preferentially interacts with upstream or with downstream regions. A bin can take on one of three states: upstream-biased, downstream-biased, or not biased, with directionally biased bins becoming TAD boundaries. TADs were called using the directionality index method implementation in TADtool [75], version as of April 23, 2018.

### Armatus

Armatus uses dynamic programming to find subgraphs in a network where the nodes are the genomic regions, and the edge weights are the interaction counts. The objective is to find the set of dense sub-graphs; subgraph density is defined as the ratio of the sum of edge weights to the number of nodes within the subgraph. Armatus version 2.3 was used for comparison.

### Insulation score

In the insulation score method, each bin is assigned an insulation score, calculated as the mean of the interaction counts in the window (of a predefined size) centered on the given bin. Bins corresponding to the local minima in the vector formed by these insulation scores are treated as TAD boundaries. TADtool [75] implementation of insulation score method, version as of April 23, 2018, was used in our experiments.

### 3DNetMod

3DNetMod employs a Louvain-like algorithm to partition a network of genomic regions into communities where the edge weights in the network are the interaction counts. It uses greedy dynamic programming to maximize modularity, a metric of network structure measuring the density of intra-community edges compared to random distribution of links between nodes. Version 1.0 (10/06/17) was used in our comparison.

### rGMAP

rGMAP trains a two-component Gaussian mixture model to group interactions into intra-domain or inter-domain contacts. Putative TAD boundary bins are identified by those with significantly higher intra-domain counts in its upstream window or downstream window of predefined size. The chromosome is then segmented into TADs flanked by these boundaries. Version as of April 23, 2018 was used for comparison.

### HiCseg

HiCseg treats the Hi-C matrix as a 2D image to be segmented, with each block-diagonal segment corresponding to a TAD. The counts within each block are modeled to be drawn from a certain distribution (e.g. Gaussian distribution for normalized Hi-C data). Using dynamic programming, HiCseg finds a set of block boundaries that would maximize the log likelihood of counts in each block being drawn from an estimated distribution. Version 1.1 was used in our experiments.

### TopDom

TopDom generates a score for each bin along the chromosome, where the score is the mean interaction count between the given bin and a set of upstream and downstream neighbors (neighborhood size is a user-specified parameter). Putative TAD boundaries are picked from a set of bins whose score forms a local minimum; false positive boundaries are filtered out with a significance test. Version 0.0.2 was used in our analysis.

### TAD evaluation criteria

We evaluated the quality of TADs using different enrichment metrics as well as internal validation metrics used for comparing clustering algorithms.

### Enrichment analysis

#### Enrichment of known architectural proteins

We estimated the enrichment of three known architectural proteins (CTCF, RAD21 and SMC3) in the TAD boundaries of five cell lines from Rao et al [36]. TAD boundaries are defined by the starting bin and the ending bin of each predicted TAD, along with one preceding the starting bin and one following the ending bin. Let *N* be the total number of bins in a chromosome, *n*_BIND_ be the number of bins with one or more ChIP-seq peaks or accessible motif sites, *n*_TAD_ be the number of TAD boundary bins, and *n*_TAD-BIND_ be the number of TAD-boundary bins with a binding event (ChIP-seq peak or accessible motif match site). The fold enrichment for a particular protein is calculated a 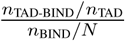 Within each cell line, the fold enrichment across all chromosomes was averaged; then the mean across cell lines was used to rank the TAD-calling methods (**Table S1**F, Supplementary Data).

### Histone modification enrichment

We used the percentage of TADs enriched in histone modification signals as a validation metric to assess the quality of TADs, similar to Zufferey et al., [28]. For each TAD, the mean histone modification ChIP-seq signal was calculated for the regions within the TAD. Next, for each TAD-calling method, TADs and non-TAD stretches were shuffled within each chromosome 10 times to yield randomized TADs. The empirical p-value of a TAD was calculated as the proportion of randomized TADs with higher mean ChIP-seq signal than that of the given TAD. A TAD was considered significantly enriched if its p-value was less than 0.05. The mean proportion of TADs with significant enrichment across cell lines was used to rank the TAD-calling methods (**Table S1**G, Supplementary Data).

### Internal validation metrics

Since a TAD represents a cluster of contiguous regions that tend to interact more among each other than with regions from another TAD or cluster, we used two internal validation or cluster quality metrics, Davies-Bouldin Index and Delta count, to evaluate the similarity of interaction profiles among regions within a TAD.

#### Davies-Bouldin Index (DBI)

The DBI for a single cluster *C*_*i*_ is defined as its similarity to its closest cluster *C*_*j*_, where *i, j* ∈ {1, …, *k*}, *i* ≠ *j*: DBI_*i*_ = max_*i*≠*j*_ *S*_*ij*_. The similarity metric, *S*_*ij*_, between *C*_*i*_ and *C*_*j*_ is defined as:

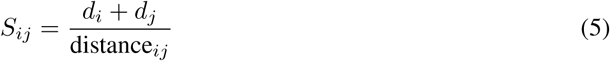

where *d*_*i*_ is the average distance between each data point in cluster *C*_*i*_ and the cluster centroid and distance_*ij*_ is the distance between the cluster centroids of *C*_*i*_ and *C*_*j*_. In applying DBI to Hi-C data, a data point consists of a vector of a genomic region’s interaction counts with other regions in the chromosome (e.g. an entire row or column in the Hi-C matrix); a cluster corresponds to a group of regions within the same TAD; the cluster centroid is a mean vector of rows that belong to the same cluster/TAD. The smaller the DBI, the more distinct the clusters are from one another.

For each TAD-calling method, we first computed the DBI for each TAD. Next, TADs and non-TAD stretches were shuffled within each chromosome 10 times to yield randomized TADs. The empirical p-value of a TAD was calculated as the proportion of randomized TADs with lower DBI (recall a lower DBI means better clustering) than that of the given TAD.

#### Delta Contact Count (DCC)

DCC for cluster *C*_*i*_ is defined as follows: let in_*i*_ denote the mean interaction counts between pairs of regions that are both in *C*_*i*_, and out_*i*_ denote the mean interaction counts between pairs of regions where one region is in cluster *C*_*i*_ and the other region is not. Then DCC_*i*_ = in_*i*_ − out_*i*_.

We expect that for a good cluster, the pairs of regions within the cluster should have higher contact counts. Therefore, the higher the value of DCC, the higher the quality of the cluster. Again, a cluster corresponds to a group of regions within the same TAD. Given the DCC values for each TAD, the empirical p-value of a TAD was calculated as the proportion of randomized TADs with higher delta count than that of the given TAD.

A TAD was considered to have significant DBI or DCC if its p-value was less than 0.05. The mean proportion of TADs with significant DBI/DCC across cell lines was used to rank the TAD-calling methods (**Table S1**A,B, Supplementary Data).

### TAD similarity and stability metrics

When assessing the similarity or stability of TADs, we used cluster comparison metrics, Rand Index and Mutual Information. First, TADs were converted to clusters so that regions in the same TAD were all assigned to the same cluster; all non-TAD regions, if a TAD-calling algorithm should have them, were assigned to a single cluster together.

For Rand Index, each genomic region is treated as a node in a graph; two nodes are connected by an edge if they are in the same cluster. Then, the number of edges that were preserved between clustering result *A* and clustering result *B* is divided by the total number of pairs of nodes, i.e. number of edges in a fully connected graph. Rand Index of 1 corresponds to perfect concordance between two clustering results; Rand Index of 0 means no agreement.

Mutual Information (MI) is an information-theoretic metric measuring the dependency between two random variables, where each variable can be a clustering result. Specifically,

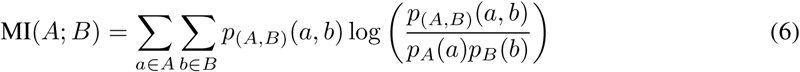

where *A, B* are random variables derived from clustering results, e.g. *A* is the cluster assignment corresponding to TADs from high-depth data and *B* is the cluster assignment based on TADs from down-sampled data. Mutual Information is 0 if the joint distribution of *A* and *B* equals the product of each marginal distribution, i.e. *A* and *B* are independent, or in an information-theoretic sense, knowing *A* does not provide any information about *B*. The higher the Mutual Information value, the greater the information conveyed by the variables about each other; in the context of measuring clustering agreement, one clustering result is similar to the other.

Both metrics were used to evaluate the stability of TADs across depth, the similarity of TADs from different TAD-calling methods, the recovery of TADs from smoothed Hi-C data, the similarity of TADs along time-course data, and the consistency of GRiNCH TADs from different 3D genome capturing technologies (e.g. SPRITE, HiChIP). In ranking TAD-calling methods for stability across depth, the mean Rand Index or Mutual Information across cell lines was used (**Table S1**D,E, Supplementary Data).

### Robustness to low-depth data

To assess the robustness or stability of TADs to low-depth input data, the TADs from a high-depth dataset (GM12878) [36] were compared to the TADs from a downsampled, low-depth dataset. If the original set of TADs are similar to the set of TADs from downsampled data, they are considered to be stable to low depth. The similarity metrics used are described in the “TAD similarity and stability metrics” section.

In order to downsample a high-depth Hi-C matrix (e.g. from GM12878) to similar levels as a lower depth one (e.g. from HMEC), a distance-stratified approach was used to match both the mean of non-zero counts and sparsity level between the two datasets. First, for each distance threshold *d*, let 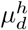 denote the mean of the non-zero counts in the high-depth dataset and 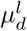 denote the mean on non-zero counts in the low-depth dataset. The scaled down value for each non-zero entry of the original high-depth dataset is: 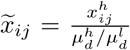 where 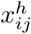 is the value for the *i,j* bin pair in the high-depth dataset. Then, to increase the sparsity of the high-depth dataset, *z*_*d*_ of the non-zero counts in the high-depth dataset at distance *d* is randomly set to zero, where *z*_*d*_ is the number of additional entries in the low-depth dataset that are zero compared to the high-depth dataset.

### Identification of novel factor enrichment at GRiNCH TAD boundaries

A similar procedure to CTCF boundary enrichment was used to identify novel boundary elements, by assessing whether the accessible motif sites of 746 transcription factors from the JASPAR core vertebrate collection [43] are enriched in GRiNCH TAD boundaries. This procedure was applied to the five cell lines from Rao et al [36] and the cell types or time points from mouse reprogramming data [42]. One change to the procedure was that instead of calculating fold enrichment per chromosome, all counts were aggregated across all chromosomes within the given cell line, cell type, or time point. The hypergeometric test was used to calculate the significance of the number of TF sites in the boundaries and were ranked based on their p-value.

### Smoothing methods

#### Smoothing with GRiNCH via matrix completion

GRiNCH smooths a noisy input Hi-C matrix by using the matrix completion aspect of NMF. Specifically, the reconstructed matrix X^*s*^ = UV^T^ is the smoothed matrix. The effectiveness of GRiNCH matrix completion as a smoothing method was compared to that of mean filter and Gaussian filter, two methods used in image blurring [76] and Hi-C datasets [30].

#### Mean filter

Mean filtering is used in HiCRep [30] as a preprocessing step to measure reproducibility of Hi-C datasets. To create a smoothed matrix X^*s*^ from a given input matrix X with a mean filter, each element in 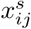 is estimated from the mean of its neighboring elements within radius 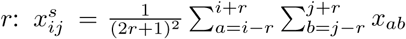. Three different values for the radius *r* were considered: *r* ∈ {3, 6, 11}.

#### Gaussian filter

A Gaussian filter uses a weighted mean of the neighborhood of a particular contact count entry, *x*_*ij*_, where the weight is determined by the distance of the neighbor from the given position:

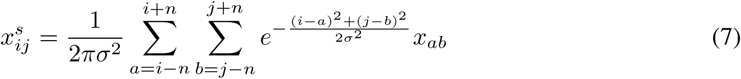

Three different values of (*σ*) were considered, *σ* ∈ {1, 2, 3} and *n* was set to 4 ∗ *σ*.

### Assessment of benefits from smoothing

#### Recovery of TADs from smoothed downsampled data

To assess whether smoothing helps preserve or recover structure from low-depth data, downsampled datasets (see “Robustness to low-depth data”) were smoothed with methods described above (see “Smoothing methods”). The Directionality Index (DI) TAD finding method was applied to the high and low depth datasets. Then the similarity of the TADs from the original high depth and the TADs from the smoothed data were measured (see “TAD similarity and stability metrics”). Higher similarity metric values imply better recovery of structure from smoothing.

#### Recovery of significant interactions using Fit-Hi-C

Fit-Hi-C [40] was used to call significant interactions in the original and the smoothed Hi-C datasets. Interactions from the original high-depth Hi-C dataset with Fit-Hi-C q-value *<* 0.05 was defined as the set of “true” significant interactions. From the downsampled then smoothed matrices, each smoothed interaction count was assigned a “prediction score” of 1− its Fit-Hi-C q-value. Precision and recall curves were then computed using the “true” interactions and the “prediction scores.” The recovery of significant interactions was measured with the Area under the Precision-Recall curve (AUPR).

## Implementation and availability

Source code (implemented in C++), installation instructions (supported in Linux distributions), documentation, and tutorial for visualization (scripts implemented in Python) can be found at roy-lab.github.io/grinch. Scripts used to analyze the results and generate the figures are available to download.

## Supporting information

Data and scripts to generate Table 1 and Table S1

## Author contributions

Lee and Roy conceptualized the overall framework and algorithm. Lee implemented the algorithm, designed and performed experiments, and wrote the manuscript. Roy designed the experiments and wrote the manuscript.

## Acknowledgements

We thank Shilu Zhang and Alireza Fotuhi Siahpirani for providing scripts for data processing and interpretation of results. We also thank the Center for High Throughput Computing at UW Madison for computational resources.

