## Supplementary material for "Simultaneous smoothing and detection of topological units of genome organization from sparse chromatin contact count matrices with matrix factorization": Figures S1-S7

### Supplementary Figure 1

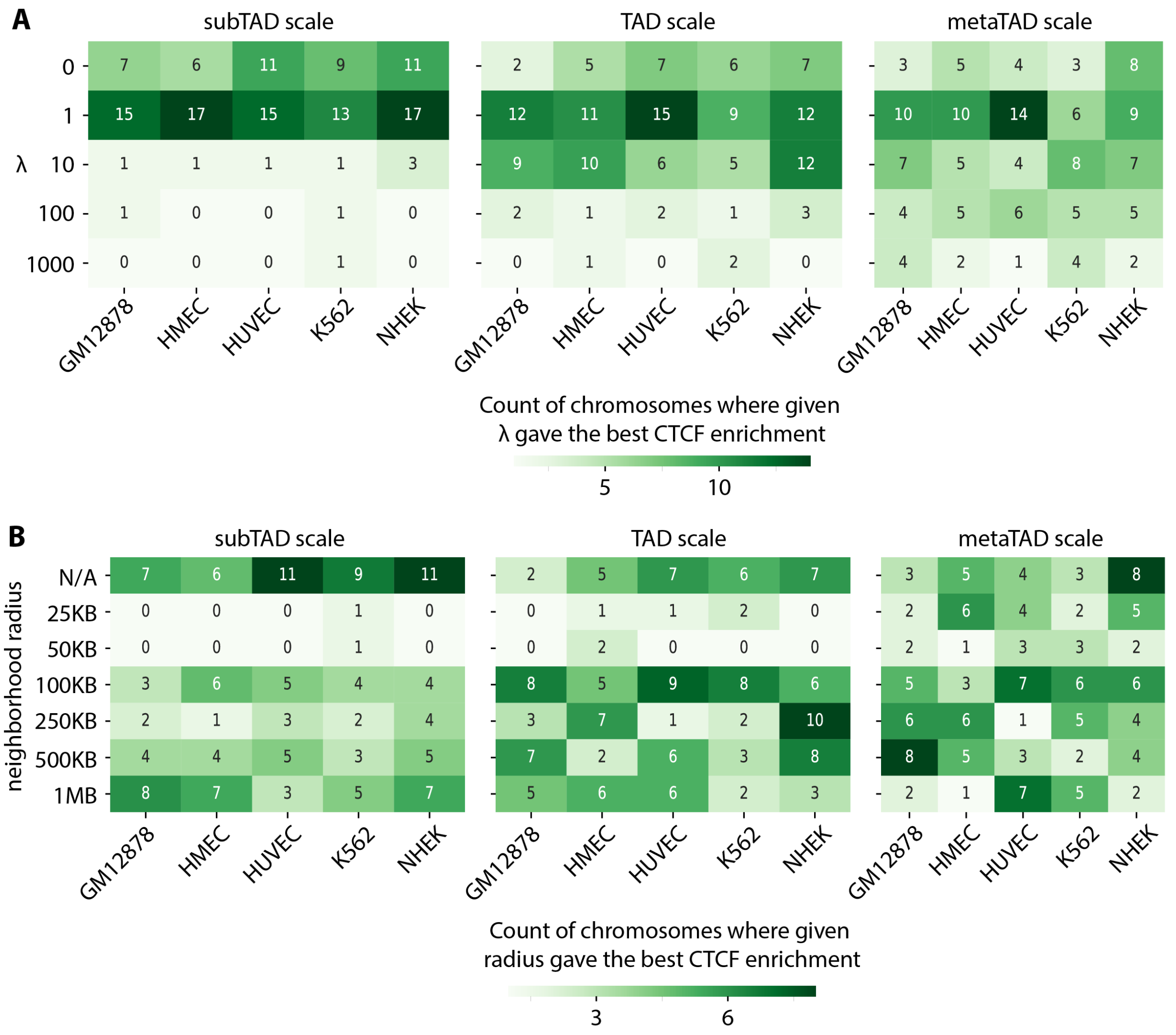

**Figure S1:** Selecting graph regularization parameters $\lambda$ and neighborhood radius, $r$. $\lambda$ controls the strength of regularization. $r$ determines how many neighboring genomic regions will be used to influence a given region during the regularization process. **A**. Shown are the count of chromosomes in which the given $\lambda$ value (row) gave the best CTCF enrichment within the given cell line (column), for different $k$ settings denoted by subTAD, TAD, and metaTAD scale (see **Figure S5**, **Figure S6**). Within each cell line, for each chromosome, we ranked the tested parameter combinations of $\lambda$ and $r$, and then counted the times a given $\lambda$ yielded the best CTCF enrichment (regardless of $r$ value). Due to ties in ranking, each column can add up to 23 or more. $\lambda = 0$ corresponds to vanilla NMF without regularization. **B**. Shown are the count of chromomes in which the given neighborhood radius value (row) gave the best CTCF enrichment in each cell line (column) at different $k$ settings. As in **A**, we rank the parameter combinations of $\lambda$ and $r$ based on the CTCF enrichment, then count the number of times a particular value of $r$ yielded the best fold enrichment. N/A corresponds to vanilla NMF without regularization.

### Supplementary Figure 2

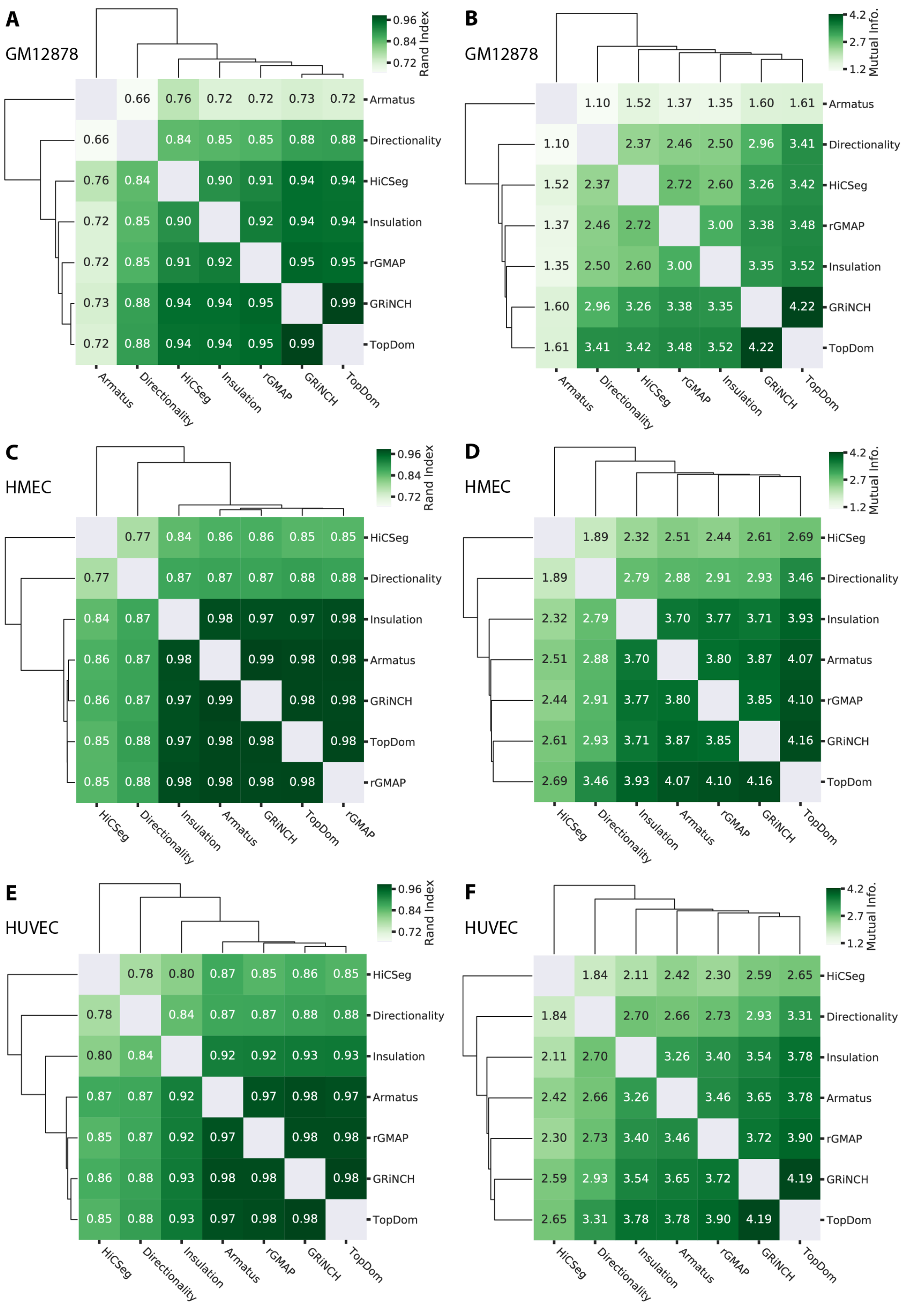

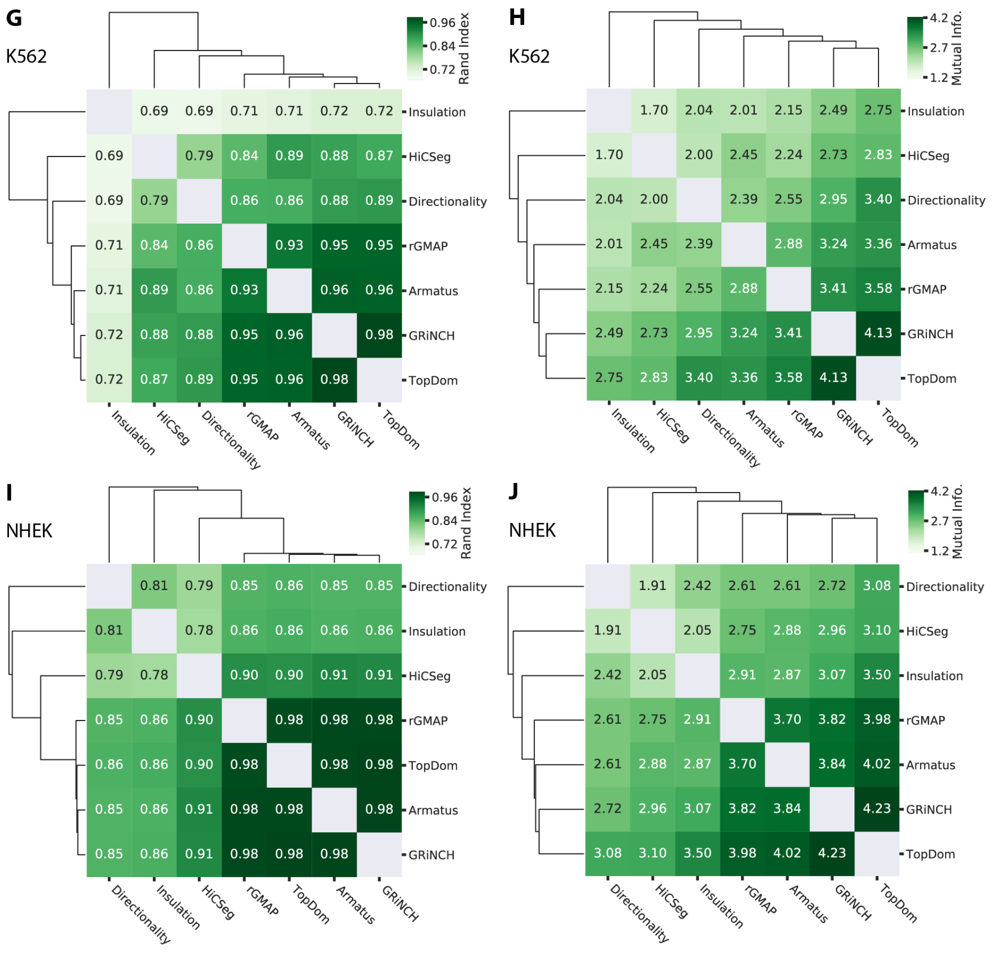

**Figure S2:** Evaluating similarity of TADs from different TAD-calling methods using two cluster quality metrics, Rand Index (**A**, **C**, **E**,**G**, **I**) and Mutual Information (**B**, **D**, **F**,**H**, **J**) for five cell lines from Rao et al., Gm12878 (**A**,**B**), HMEC (**C**,**D**), HUVEC (**E**,**F**), K562 (**G**, **H**), NHEK (**I**,**J**).

### Supplementary Figure 3

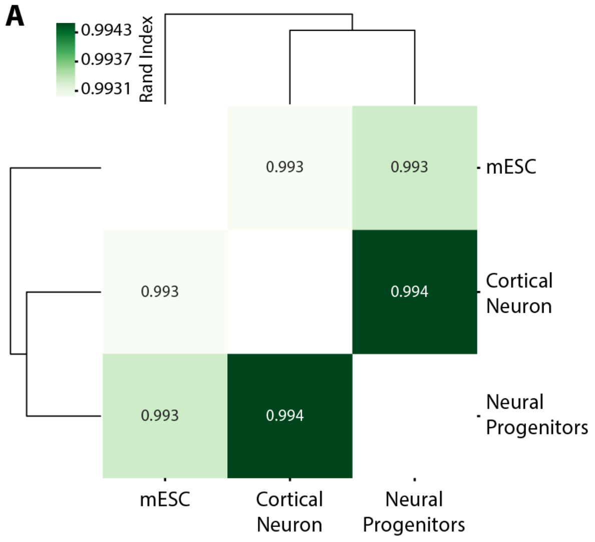

**Figure S3:** Similarity of GRiNCH TADs from mouse neural development time-course Hi-C data measured for three stages: mESC, Cortical Neuron, Neural Progenitors. The order of the developmental stages is mESC, Cortical Neurons, Neural Progenitors. **A**. Similarity of TADs by measured by Rand index. **B**. Similarity of TADs measured by Mutual Information.

### Supplementary Figure 4

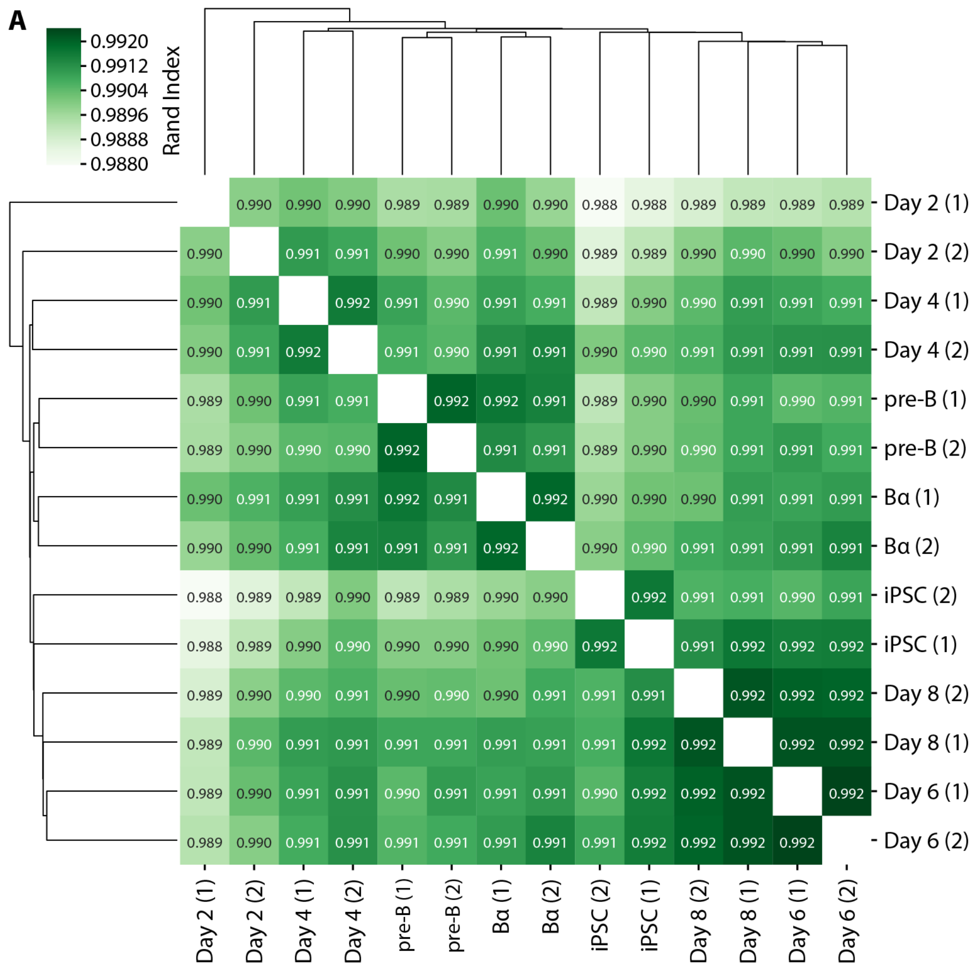

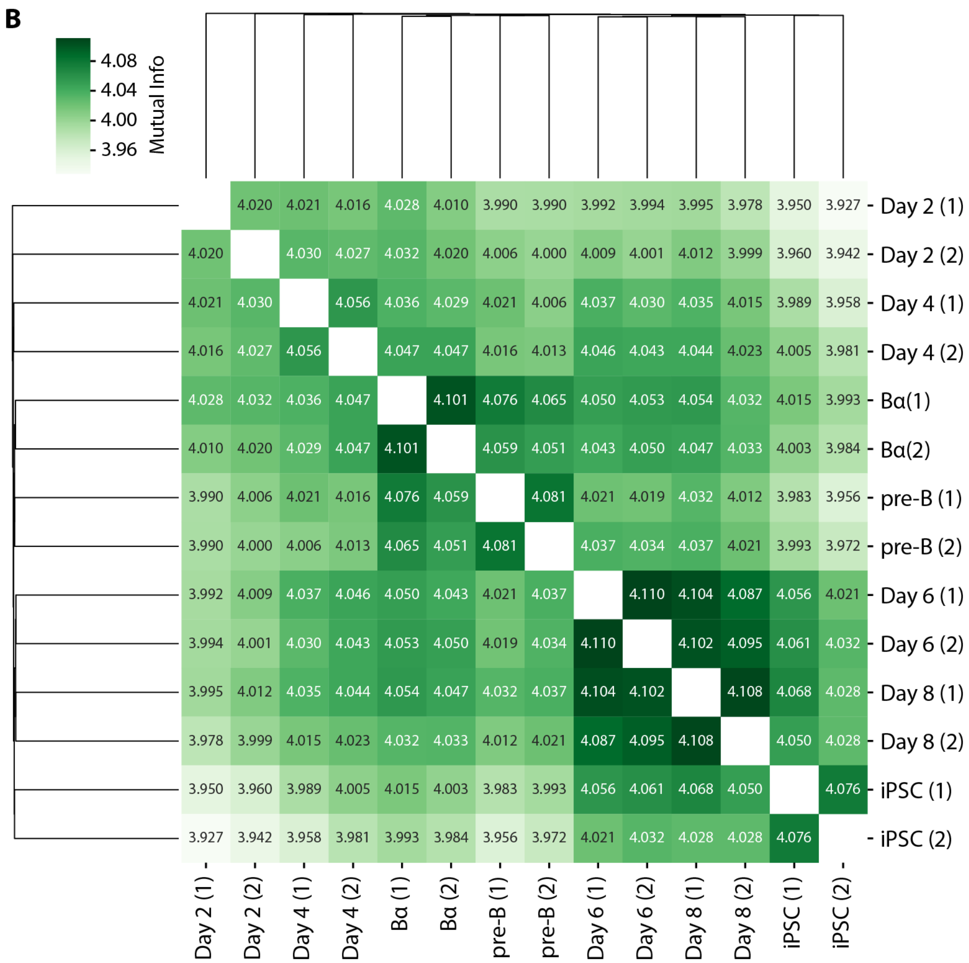

**Figure S4:** Similarity of GRiNCH TADs from pluripotency reprogramming time-course Hi-C data measured for the starting pre-B cells and ending induced pluripotent (iPSC) state and five intermediate time points, B$\alpha$, Day2, Day4, Day6, Day8. (1) and (2) suffixes represent replicate 1 and 2 respectively. **A**. Similarity of TADs measured by Rand index. **B**. Similarity of TADs measured by Mutual Information.

### Supplementary Figure 5

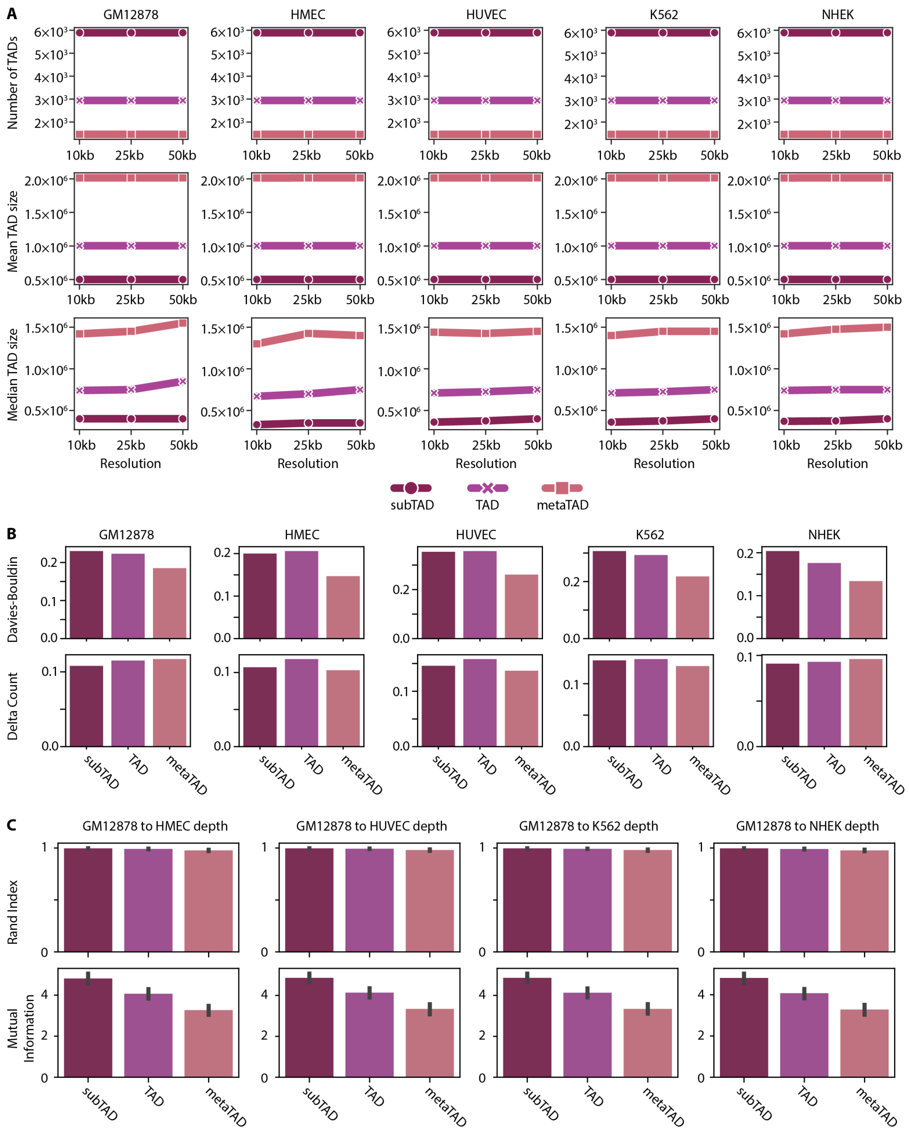

**Figure S5:** Shown are different statistics for different settings of the number of clusters in GRiNCH, $k$. We set $k$ based on the expected size of the clusters and consider three expected sizes: subTADs (500kb), TADs (1MB), metaTADs (2MB). TADs are known to be ∼1MB and therefore, for the “TAD scale” regions, $k = \frac{n_c}{1\text{Mb}}$, where $n_c$ is the length of chromosome $c$. **A**. The number of subTADs, TADs, and metaTADs, and their median size for Hi-C datasets from five cells lines at three different resolutions: 10kb, 25kb, 50kb. **B**. Proportion of subTADs, TADs, or metaTADs with significantly better (p-val $< 0.05$) cluster quality metrics than random clusters, as measured by Davies-Bouldin Index and Delta Count. Results here are shown for 25kb-resolution data. **C**. The similarity between subTADs, TADs, and metaTADs from high-depth GM12878 dataset and those from datasets downsampled to other lower depths available in different cell lines. Similarity is measured by Rand Index and Mutual Information. Results here are shown for 25kb-resolution data.

### Supplementary Figure 6

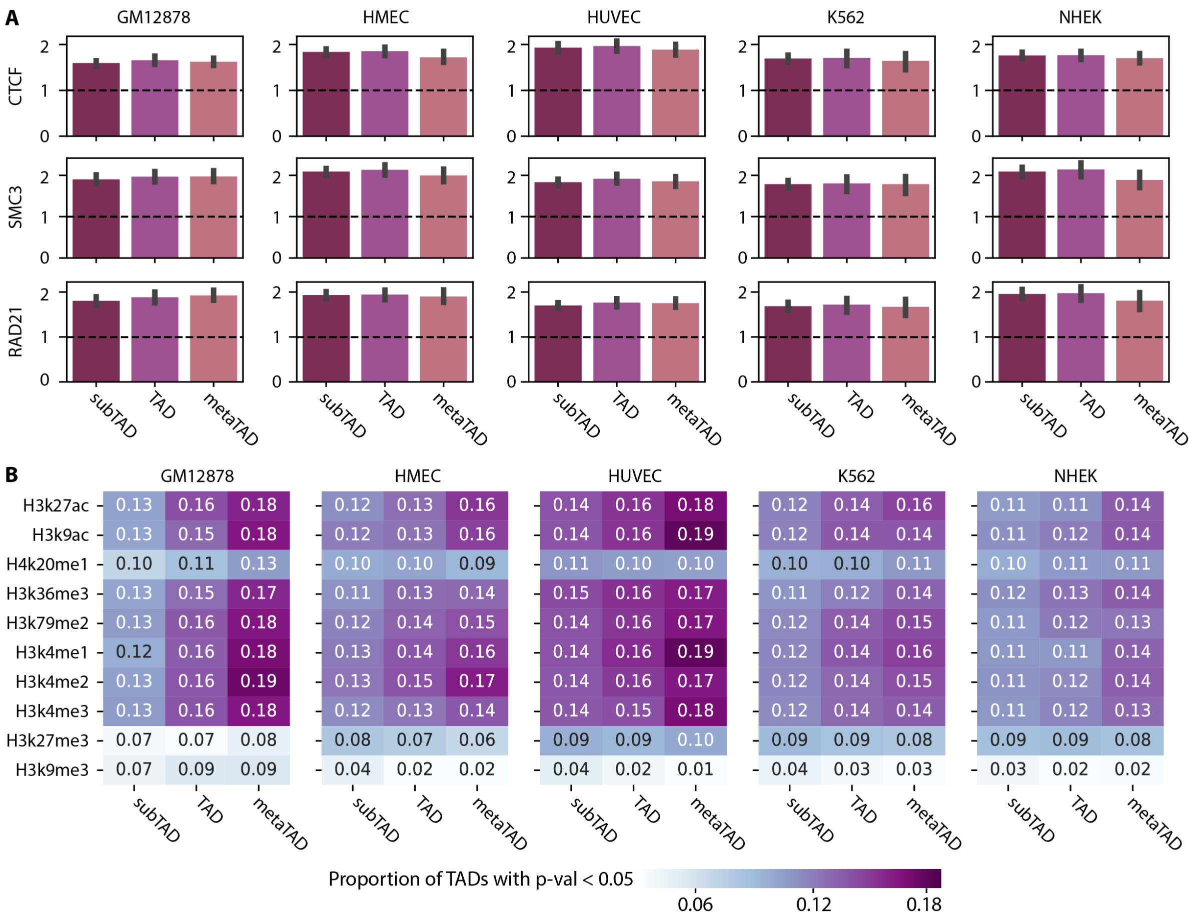

**Figure S6:** Enrichment of regulatory signals in GRiNCH clusters of different expected sizes: subTAD (500kb), TAD (1Mb) , and metaTAD (2Mb). The expected size is used to set the GRiNCH parameter $k$, the number of clusters. Results are shown for 25kb resolution data. **A**. Fold enrichment of architectural protein binding signals in subTAD, TAD, and metaTAD boundaries. **B**. Proportion of subTADs, TADs, and metaTADs with significantly higher values (p-val $<$ 0.05) of mean histone modifications compared to random clusters.

### Supplementary Figure 7

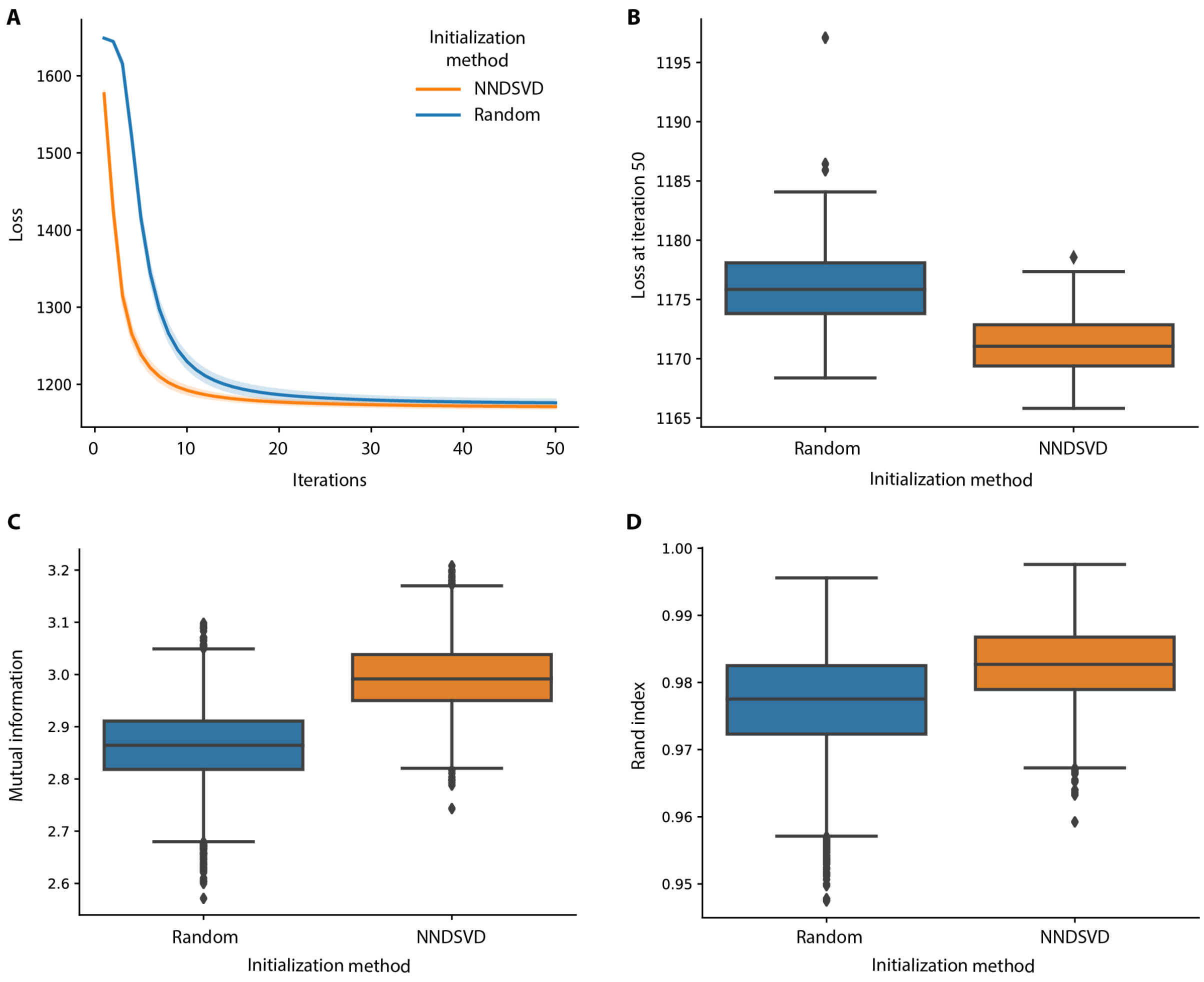

**Figure S7:** Comparison of NNDSVD initialization versus random initialization. **A**. Loss as a function of iterations of the GRiNCH algorithm with NNDSVD initialization versus random initialization. For each type of initialization, we tracked the value of the objective/loss over 50 iterations of the GRiNCH algorithm, using 100 different seeds for randomization. The solid line represent the mean loss at a given iteration, and the lightly colored bands around each line is the standard deviation of the loss at the given iteration. **B**. The distribution of the loss after 50 iterations for each type of initialization. Each point on the box plot corresponds to one of 100 different random seeds. **C**. Measuring stability of GRiNCH TADs from NNDSVD initialization versus random initialization. Each box plot shows the distribution of similarity score of GRiNCH TADs using different initialization methods. GRiNCH TADs after 50 iterations from each run with different seeds and initialization methods were converted to clusters. Pairwise similarity of clustering results from seed $x$ and seed $y$ using the same initialization method was measured with mutual information. Higher mutual information means the GRiNCH TADs tend to be more similar and therefore more stable across different seeds. **D**. Measuring stability of GRiNCH TADs with Rand Index. GRiNCH TADs were generated in the same as in **C**. Higher Rand index means the GRiNCH TADs tend to be more similar and therefore more stable across different seeds.

### List of Supplementary Tables

**Table S1(A)** Ranking TAD-calling methods by mean proportion of TADs with significant Davies-Bouldin Index across 5 cell lines

**Table S1(B)** Ranking TAD-calling methods by mean proportion of TADs with significant Delta Contact Count across 5 cell lines

**Table S1(C)** Ranking TAD-calling methods by mean (absolute) change in median TAD size as input data resolution changes from 10kb to 25kb to 50kb

**Table S1(D)** Ranking TAD-calling methods by mean Rand Index between TADs from high-depth Gm12878 data and TADs from Gm12878 data downsampled to other cell lines' depth.

**Table S1(E)** Ranking TAD-calling methods by mean mutual information between TADs from high-depth Gm12878 data and TADs from Gm12878 data downsampled to other cell lines' depth.

**Table S1(F)** Ranking TAD-calling methods by mean fold enrichment of known boundary elements (CTCF, SMC3, RAD21) across 5 cell lines.

**Table S1(G)** Ranking TAD-calling methods by mean proportion of TADs with significant histone modification signals across 5 cell lines

**Table S2** Ranking of transcription factors by significant motif enrichment in TAD boundaries across all cell types or time points during mouse pluripotency reprogramming (mouse pre-B cell, D2, D4, D6, D8, iPSC)

**Table S3** Ranking of transcription factors by significant motif enrichment in TAD boundaries across all cell lines (GM12878, HUVEC, HMEC, NHEK, K562) from Rao et al.
