## Supplementary material for "Simultaneous smoothing and detection of topological units of genome organization from sparse chromatin contact count matrices with matrix factorization": Data and scripts to generate Table 1 and Table S1: deltacount_prop_pivot.py

```
"""
This script ranks TAD calling methods by the average proportion of TADs with significant delta count
across 5 cell lines.
input: deltacount_prop.txt
usage: python deltacount_prop_pivot.py
output: deltacount_prop_pivot.txt
"""
import pandas as pd

df = pd.read_csv('deltacount_prop.txt',sep="\t")
df = df.loc[(df.Method != 'GRiNCH-subTAD') & (df.Method != 'GRiNCH-metaTAD')]
df_pivot = df.pivot('Method','Cell','p-val<0.05')
df_pivot = df_pivot.rename_axis(None,axis=1).reset_index()
df_pivot.set_index('Method',inplace=True)
df_pivot['Mean across cell lines'] = df_pivot.mean(axis=1,numeric_only=True)
df_pivot['Rank by mean'] = df_pivot['Mean across cell lines'].rank(method='dense', ascending=False).astype(int)
df_pivot.to_csv('deltacount_prop_pivot.txt',sep="\t", float_format="%.3f")
```
