## Supplementary material for "Simultaneous smoothing and detection of topological units of genome organization from sparse chromatin contact count matrices with matrix factorization": Data and scripts to generate Table 1 and Table S1: tad_median_size_pivot.py

```
"""
This script ranks TAD calling methods by average (absolute) change in median TAD size going from 10kb input data resolution to 50kb.
input: tad_count_and_size.txt
usage: python tad_median_size_pivot.py
output: tad_median_size_pivot.txt
"""
import pandas as pd

df = pd.read_csv('tad_count_and_size.txt',sep="\t")
df = df.loc[df['Number of TADs'] > 0]
df['Cell-res'] = df.Cell + '-' + df.Resolution

df_pivot = df.pivot('Method','Cell-res','Median TAD Size')
df_pivot = df_pivot.rename_axis(None,axis=1).reset_index()
df_pivot.set_index('Method',inplace=True)
for cell in ['Gm12878','Hmec','Huvec','K562','Nhek']:
	df_pivot[cell + ' 10kb->25kb'] = (df_pivot[cell+'-10kb'] - df_pivot[cell+'-25kb']).abs()
	df_pivot[cell + ' 25kb->50kb'] = (df_pivot[cell+'-50kb'] - df_pivot[cell+'-25kb']).abs()
	df_pivot.drop(columns=[cell+'-10kb',cell+'-25kb',cell+'-50kb'],inplace=True)

df_pivot['Mean across cell lines'] = df_pivot.mean(axis=1,numeric_only=True)
df_pivot['Rank by mean'] = df_pivot['Mean across cell lines'].rank(method='dense', ascending=True).astype(int)
df_pivot.to_csv('tad_median_size_pivot.txt',sep="\t", float_format="%.3f")
```
