## Supplementary material for "Simultaneous smoothing and detection of topological units of genome organization from sparse chromatin contact count matrices with matrix factorization": Data and scripts to generate Table 1 and Table S1: mutualinfo_mean_pivot.py

```
"""
This script ranks TAD calling methods by the average mutual information across 5 different depths of downsampling.
input: compare.txt
usage: python mutualinfo_mean_pivot.py
output: mutualinfo_mean_pivot.txt
"""
import pandas as pd

df = pd.read_csv('compare.txt',sep="\t",usecols=['Method','Depth','MI'])
df = df.loc[(df.Method != 'GRiNCH-subTAD') & (df.Method != 'GRiNCH-metaTAD')]
df = df.groupby(['Method', 'Depth']).mean().reset_index()
df_pivot = df.pivot('Method','Depth','MI')
df_pivot = df_pivot.rename_axis(None,axis=1).reset_index()
df_pivot.set_index('Method',inplace=True)
df_pivot['Mean across depths'] = df_pivot.mean(axis=1,numeric_only=True)
df_pivot['Rank by mean'] = df_pivot['Mean across depths'].rank(method='dense', ascending=False).astype(int)
df_pivot.to_csv('mutualinfo_mean_pivot.txt',sep="\t", float_format="%.3f")
```
