## Supplementary material for "Simultaneous smoothing and detection of topological units of genome organization from sparse chromatin contact count matrices with matrix factorization": Data and scripts to generate Table 1 and Table S1: fold_enrichment_mean_pivot.py

```
"""
This script ranks TAD calling methods by the average boundary element fold enrichment across 5 different cell lines.
input: boundary_enrichment.txt
usage: python fold_enrichment_mean_pivot.py
output: fold_enrichment_mean_pivot.txt
"""
import pandas as pd

df = pd.read_csv('boundary_enrichment.txt',sep="\t")
df = df.loc[(df.Method != 'GRiNCH-subTAD') & (df.Method != 'GRiNCH-metaTAD')]
df[['Cell','Chro']] = df['Cell-chro'].str.split('-',expand=True)
df[['TF','type']] = df['Signal'].str.split(' ',expand=True)
df['Cell-TF'] = df.Cell +'-'+df.TF
df = df[['Cell-TF','Method','Chro','Fold Enrichment']]
df = df.groupby(['Method', 'Cell-TF']).mean().reset_index()
df_pivot = df.pivot('Method','Cell-TF','Fold Enrichment')
df_pivot = df_pivot.rename_axis(None,axis=1).reset_index()
df_pivot.set_index('Method',inplace=True)
df_pivot['Mean across cell lines'] = df_pivot.mean(axis=1,numeric_only=True)
df_pivot['Rank by mean'] = df_pivot['Mean across cell lines'].rank(method='dense', ascending=False).astype(int)
df_pivot.to_csv('fold_enrichment_mean_pivot.txt',sep="\t", float_format="%.3f")
```
