## Supplementary material for "Simultaneous smoothing and detection of topological units of genome organization from sparse chromatin contact count matrices with matrix factorization": Data and scripts to generate Table 1 and Table S1: README

Each directory contains the per-chromosome-level data used to rank TAD-calling methods for different criteria. 

Directory A contains the data for Table S1(A), so on and so forth. 

In each directory, X_pivot.txt contains the aggregated data with ranking, generated by X_pivot.py script. The other text file contains the per-chromosome-level data before aggregation. See the header of X_pivot.py file for usage and the input, output files.
